# SARS-CoV-2 Airway Infection Results in Time-dependent Sensory Abnormalities in a Hamster Model

**DOI:** 10.1101/2022.08.19.504551

**Authors:** Randal A. Serafini, Justin J. Frere, Jeffrey Zimering, Ilinca M. Giosan, Kerri D. Pryce, Ilona Golynker, Maryline Panis, Anne Ruiz, Benjamin tenOever, Venetia Zachariou

## Abstract

Despite being largely confined to the airways, SARS-CoV-2 infection has been associated with sensory abnormalities that manifest in both acute and long-lasting phenotypes. To gain insight on the molecular basis of these sensory abnormalities, we used the golden hamster infection model to characterize the effects of SARS-CoV-2 versus Influenza A virus (IAV) infection on the sensory nervous system. Efforts to detect the presence of virus in the cervical/thoracic spinal cord and dorsal root ganglia (DRGs) demonstrated detectable levels of SARS-CoV-2 by quantitative PCR and RNAscope uniquely within the first 24 hours of infection. SARS-CoV-2-infected hamsters demonstrated mechanical hypersensitivity during acute infection; intriguingly, this hypersensitivity was milder, but prolonged when compared to IAV-infected hamsters. RNA sequencing (RNA-seq) of thoracic DRGs from acute infection revealed predominantly neuron-biased signaling perturbations in SARS-CoV-2-infected animals as opposed to type I interferon signaling in tissue derived from IAV-infected animals. RNA-seq of 31dpi thoracic DRGs from SARS-CoV-2-infected animals highlighted a uniquely neuropathic transcriptomic landscape, which was consistent with substantial SARS-CoV-2-specific mechanical hypersensitivity at 28dpi. Ontology analysis of 1, 4, and 30dpi RNA-seq revealed novel targets for pain management, such as ILF3. Meta-analysis of all SARS-CoV-2 RNA-seq timepoints against preclinical pain model datasets highlighted both conserved and unique pro-nociceptive gene expression changes following infection. Overall, this work elucidates novel transcriptomic signatures triggered by SARS-CoV-2 that may underlie both short- and long-term sensory abnormalities while also highlighting several therapeutic targets for alleviation of infection-induced hypersensitivity.

**One Sentence Summary:** SARS-CoV-2 infection results in an interferon-associated transcriptional response in sensory tissues underlying time-dependent hypersensitivity.

## INTRODUCTION

COVID-19, the disease resulting from SARS-CoV-2 infection, is associated with highly variable clinical outcomes that range from asymptomatic disease to death. For milder infections, COVID-19 is associated with primarily respiratory infection-associated symptoms (cough, congestion, fever) and sensory phenotypes such as headache and anosmia (*1–3*). For more severe cases, however, SARS-CoV-2 infection has been seen to induce a variety of systemic perturbations that have the ability to affect nearly every organ, including strokes from vascular occlusion, cardiovascular damage, and acute renal failure (*4–6*). Intriguingly, recent research has shown that a significant number of actively infected patients suffering from both mild and severe infections experience sensory-related symptoms such as headache, visceral pain, Guillain-Barre syndrome (GBS), nerve pain, and polyneuritis (*7–9*). While these symptoms subside after clearance of infection in a majority of patients, they have been noted to arise in or persist to sub-acute or chronic timepoints for many (*10, 11*). Sensory-related symptomology is thus a major component of long COVID, a condition defined by the World Health Organization as the persistence of COVID-19-associated symptomology that lasts for at least two months and cannot be explained by an alternative diagnosis (*12*). Accordingly, high persistence of abdominal, chest, and muscle pains, as well as headaches, was observed in long COVID patients (*13–15*). Of note, the medical field has observed a high prevalence of asymptomatic acute COVID-19 cases, suggesting a mechanistic divergence between the acute and chronic stages of the disease (*16, 17*).

Abnormal somatosensation is a common symptom of neuroinvasive and non-neuroinvasive viral infections, including varicella-zoster virus (VZV), human immunodeficiency virus (HIV), and SARS-CoV-2 (*18–20*). Phenotypes generally consist of painful sensations (burning, prickling, or aching), as well as paresthesias (tingling) or in some cases hypoesthesias (numbness). The mechanisms underlying these symptoms vary by virus. For example, certain neurotropic viruses, such as herpesviruses, persist in the dorsal root ganglia (DRGs) and directly induce abnormal activity in these primary sensory cells upon reactivation (*21, 22*).Retroviruses, such as HIV, can induce primary sensory neuropathy through viral protein interaction with axons, while also inducing secondary inflammation at these neural sites, thereby inducing hyperexcitability and chronic pain symptoms (*23, 24*). However, the mechanisms by which coronaviruses, and specifically SARS-CoV-2, induce abnormal sensation are more poorly understood (*25*).

The ability of SARS-CoV-2 to pass the blood brain barrier and directly infect the central nervous system (CNS) is currently unclear. While ultrastructural analyses of post-mortem tissue from COVID-19 patients have identified structures resembling viral particles in the central nervous system, other studies have failed to detect replication-competent virus in the brain (*26–30*). These seemingly disparate results may be explained by pre-clinical studies in the golden hamster model of SARS-CoV-2 infection, which demonstrate the presence of viral RNA in various brain regions, including the olfactory bulb, cortical areas, brainstem, and cerebellum during acute infection, despite lacking evidence for any infectious material (*31, 32*). Together, these data suggest that virus replication in the airways results in the dissemination of viral RNA and the induction of an antiviral transcriptional response in distal tissues, including the brain, which may underly CNS-related pathologies, such as demyelinating lesions and brain hemopathologies, observed among COVID-19 patients (*33–36*).

Despite the large number of studies investigating CNS infiltration during SARS-CoV-2 infection, little clinical or pre-clinical literature has investigated penetration capabilities of SARS-CoV-2 into the peripheral nervous system, particularly sensory components such as the dorsal root ganglia (DRGs) and spinal cord (SC). While several case studies have highlighted peripheral fiber neuropathies in actively and recovered COVID-19 patients (*37, 38*), there is conflicting data regarding the presence of viral transcripts in the cerebrospinal fluid of COVID-19 patients (*39–42*).

Previous work with other members of the coronavirus family, including hemagglutinating encephalomyelitis virus (HEV), has identified active replication and satellite-mediated sequestration of virus in rodent DRGs (*43*). Mouse hepatitis virus also demonstrates anterior spinal cord segment invasion and persistence through neuroanatomical pathways such as the olfactory bulb and trigeminal ganglia, leading to consequences such as demyelination (*44, 45*).In these studies, the source of coronavirus-induced neurological dysfunction was linked to two major causes: direct virus-induced damage and collateral damage to host cells by anti-viral immune responses (*46, 47*). While the cause of SARS-CoV-2-linked sensory pathologies is unknown, SARS-CoV-2 has been shown to infect human neuronal cells *in vitro* (*48, 49*) and to induce a robust *in vivo* systemic inflammatory response (*31, 32*). Both findings could present causal mechanisms underlying SARS-CoV-2-linked sensory symptoms.

We hypothesized that clinically-observed sensory symptoms (both positive and negative) arise from exposure of neurons in the DRG and/or spinal cord to mature virus and/or through circulating inflammatory materials including both cytokines and pathogen-associated molecular patterns (PAMPs). In order to test this hypothesis, we used the Syrian golden hamster model of SARS-CoV-2 respiratory infection, which can accurately phenocopy COVID-19 in the absence of any virus or host adaptation (*31, 32, 50*). To this end, we used gene/protein quantification and imaging techniques to assess the presence of SARS-CoV-2 in sensory tissues at acute timepoints in cervical and thoracic levels using gene and protein quantification and imaging techniques. We also applied the Von Frey assay to characterize SARS-CoV-2-induced changes in mechanical hypersensitivity compared to mock and IAV-infected hamsters at acute and chronic timepoints. Moreover, in an effort to better define the molecular underpinnings of virus-induced changes in sensory hypersensitivity, we performed RNA sequencing (RNA-seq) of thoracic DRGs from infected hamsters at 1, 4, and 31 dpi timepoints. We also used ontological analysis of predicted upstream regulators associated with the sequencing results to identify and validate novel therapeutic targets. Lastly, we performed a meta-analysis of 1, 4, and 31 dpi SARS-CoV-2 thoracic DRG sequencing data against publicly-available DRG datasets from rodent pain models to highlight relevant injury response pathways. Our findings provide insights into the sensory-altering mechanisms induced by respiratory SARS-CoV-2 infection and have the potential to guide the development of novel therapeutics for a variety of pain conditions.

## RESULTS

### SARS-CoV-2 RNA Infiltrates Thoracic and Cervical DRG and Spinal Cord Tissue

We first sought to determine if SARS-CoV-2 genetic material is present in the sensory nervous system tissues and to investigate if this presence was associated with induction of an antiviral response. To this end, we performed a longitudinal cohort study in which hamsters were treated intranasally with SARS-CoV-2 or PBS (mock-infected). Cervical and thoracic levels of DRGs and spinal cord were harvested at 1, 4, 7, and 14dpi in both groups and assessed for the presence of SARS-CoV-2 subgenomic nucleocapsid protein (*N*) and canonical type-I interferon stimulated gene *Isg15* transcripts via quantitative reverse transcription PCR (RT-qPCR). We found a substantial elevation of *N* transcripts at 1dpi in cervical DRGs (**Figure 1A**; two-way ANOVA interaction F(3,35)=4.205, p=0.0122; multiple t-tests 1dpi t=2.698, df=13, p=0.0183), cervical SC (**Figure 1B**; two-way ANOVA interaction F(3,35)=3.809, p=0.0189; multiple t-tests 1dpi t=2.392, df=11, p=0.0358), thoracic DRGs (**Figure 1E**; two-way ANOVA interaction F(3,36)=3.812, p=0.018; multiple t-tests 1dpi, t=2.528, df=13, p=0.0252), and thoracic SC (**Figure 1F**; two-way ANOVA interaction F(3,34)=4.266, p=0.0116; multiple t-tests 1dpi t=3.068, df=11, p=0.0107). Viral RNA appeared to be cleared in most samples by 4dpi. *Isg15* mRNA levels, which are generally representative of interferon signaling (*51*), had similar elevation patterns to those of *N* in cervical DRGs (**Figure 1C**; two-way ANOVA interaction F(3,35)=3.689, p=0.0208, multiple t-tests 1dpi t=3.152, df=13, p=0.00764; 4dpi t=2.361, df=12, p=0.0360), cervical SC (**Figure 1D**; two-way ANOVA interaction F(3,35)=5.001, p=0.0054; multiple t-tests 1dpi t=6.034, df=12, p=0.0000590; 4dpi t=2.656, df=13, p=0.0198), thoracic DRGs (**Figure 1G**; two-way ANOVA interaction F(3,36)=1.856, p=0.155; multiple t-tests 1dpi t=3.541, df=13, p=0.00362; 4dpi t=2.311, df=13, p=0.0379), and thoracic SCs (**Figure 1H**; two-way ANOVA interaction F(3,35)=8.478, p=0.0002; multiple t-tests 1dpi t=4.286, df=12, p=0.00106; 4dpi t=3.333, df=13, p=0.00539).

**Figure 1.**
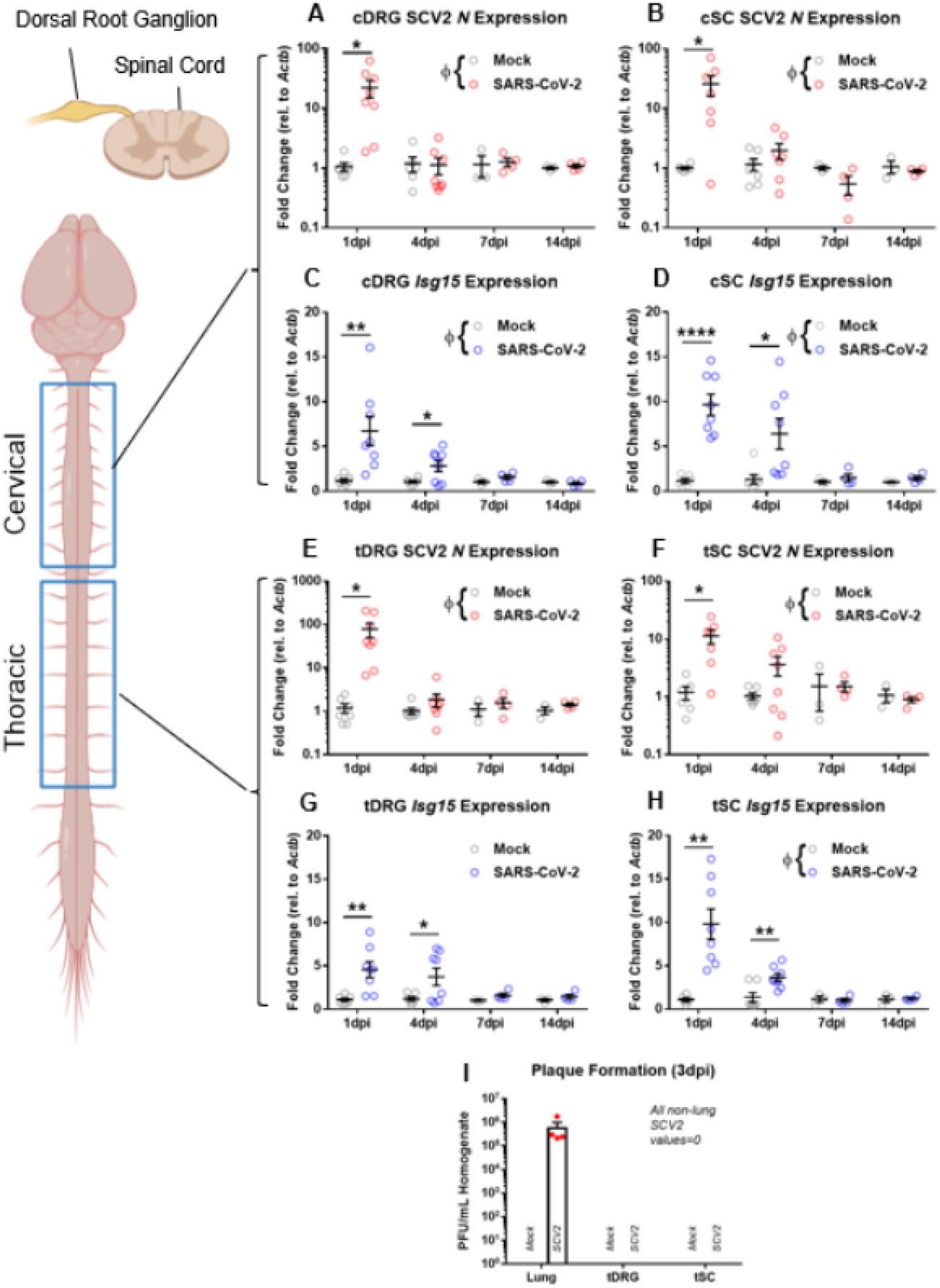
Viral mRNA and interferon-stimulated transcripts are acutely elevated in dorsal root ganglia & spinal cord by qPCR. **A-B, E-F**) Nucleocapsid protein-encoding gene (*N*) was significantly elevated in the cervical and thoracic segments of DRGs and SC at 1dpi, but not 4, 7, or 14dpi of SARS-CoV-2-infected hamsters (n=3-8/group). **C-D, G-H**) Interferon-stimulated gene 15 (*Isg15*) was significantly elevated at 1 and 4dpi at both DRG and SC levels in SARS-CoV-2-infected animals (n=3-8 per group). **I**) Plaque formation assay demonstrates mature virus presence only in the lungs of SARS-CoV-2-infected hamsters at 3dpi, but not DRG or SC (n=4 per group). Φp<0.05 for two-way ANOVA interaction factor; *p<0.05, **p<0.01 for multiple t-tests.

In order to gain insight into viral replication within the DRG, we performed a plaque assay in which combined cervical and thoracic DRGs or SC were collected at 3dpi and homogenized in PBS. This solution was then plated with Vero cells, with the number of ensuing plaques representing the number of mature virions present in the harvested tissue. As seen in **Figure 1I**, plaques were observed only in 3dpi lung homogenate from SARS-CoV-2-infected animals, but not in mock lung or any DRG or SC tissue. This suggested that mature virus was not reaching the peripheral or central sensory nervous systems.

We next sought to determine whether SARS-CoV-2 transcripts were localized to specific cell types in the DRG, which is predominantly composed of primary sensory neurons and satellite glial cells. By using RNAscope *in situ* hybridization on 1dpi cervical and thoracic cell tissue, we observed the presence of RNA (*S*) puncta around DAPI-labeled nuclei, which in DRGs are representative of satellite glial cells and *Rbfox3-labeled* neuronal spaces, but not in mock samples (**Figure 2A**). We also detected *S* transcript puncta near DAPI signal throughout SARS-CoV-2-infected cervical and thoracic spinal cord sections on 1dpi, but not in mock samples (**Figure 2B**).

**Figure 2.**
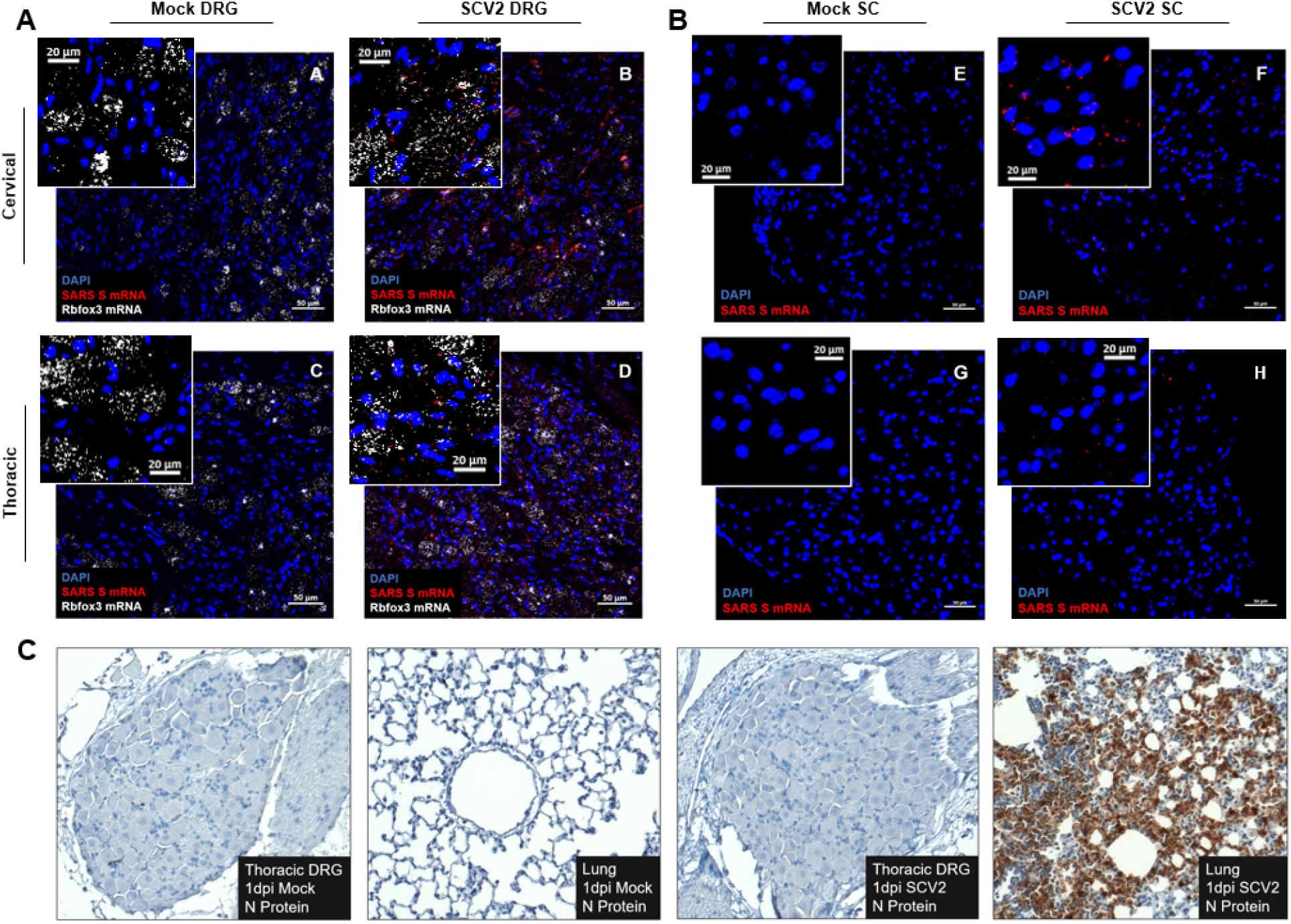
Viral mRNA is visibly detectable in DRG and SC by ISH, but mature virus is not detectable by IHC. **A**) Spike RNA (red) is detectable around DAPI (blue) and Rbox3 (white) in cervical and thoracic DRGs of SARS-CoV-2-infected animals but not mocks, suggesting infiltration of glial and neuronal cells at 1dpi (n=2/group). **B**) Spike RNA (red) is detectable around DAPI (blue) in cervical and thoracic SC, but not in mock SC at 1dpi (n=2 per group). **C**) Nucleocapsid protein was not detectable in cervical or thoracic DRGs of SARS-CoV-2-infected or mock animals, but it was detectable in lung tissue of SARS-CoV-2-infected animals at 1dpi (n=2 per group).

Of note, when tissue sections obtained from the DRGs of SVC2- or mock-infected hamsters were immuno-labeled for SARS-CoV-2 nucleocapsid protein (NP) we did not observe any notable viral protein presence (**Figure 2C**). Importantly, we confirmed the presence of NP in SARS-CoV-2-infected lung samples, but not in mock controls (**Figure 2C**). This introduced the question of whether the presence of viral mRNA and associated antiviral response signatures in the sensory nervous system are sufficient to induce behavioral and/or transcriptional perturbations.

### SARS-CoV-2 and IAV Induce Unique Mechanical Hypersensitivity Signatures

We next sought to determine whether the presence of SARS-CoV-2 RNA or associated type I interferon (IFN-I) signaling, as reported previously (*52*), was associated with the induction of sensory hypersensitivity. To assess this, we performed the Von Frey assay on hamsters infected with either IAV (A/California/04/2009) or SARS-CoV-2. IAV, similar to SARS-CoV-2, is an RNA virus of the respiratory tract that is known to provoke a systemic inflammatory response similar to SARS-CoV-2, resulting in clinically-associated myalgias (*32, 53*). Von Frey thresholds were measured during the acute phase of infection (1 and 4dpi) to identify the effects of active and subsiding SARS-CoV-2 mRNA presence and IFN-I response on sensation. As seen in **Figure 3A**, we observed a significant interaction effect between time and virus on mechanical hypersensitivity (RM two-way ANOVA Interaction F(4,18)=4.16, df=4, p=0.0147). IAV induced robust hypersensitivity at 1dpi which completely subsided by 4dpi (one-way ANOVA F=6.092, p=0.0359; Tukey’s m.c.: Baseline vs. 1dpi q=4.604, df=6, p=0.0398). SARS-CoV-2 infection instead resulted in a gradual exacerbation of hypersensitivity, reaching significance only at 4dpi (one-way ANOVA F=9.772, p=0.013; Tukey’s m.c.: Baseline vs. 4dpi q=6.117, df=6, p=0.0117). Importantly, 1dpi IAV-induced hypersensitivity was significantly higher than that caused by 1dpi SARS-CoV-2 (RM two-way ANOVA Tukey’s m.c. q=4.033, df=27, p=0.0218). Considering the emergence of distinct behavioral signatures irrespective of systemic interferon responses induced by these two viruses, we performed a time-dependent transcriptional comparison of sensory structures after infection.

**Figure 3.**
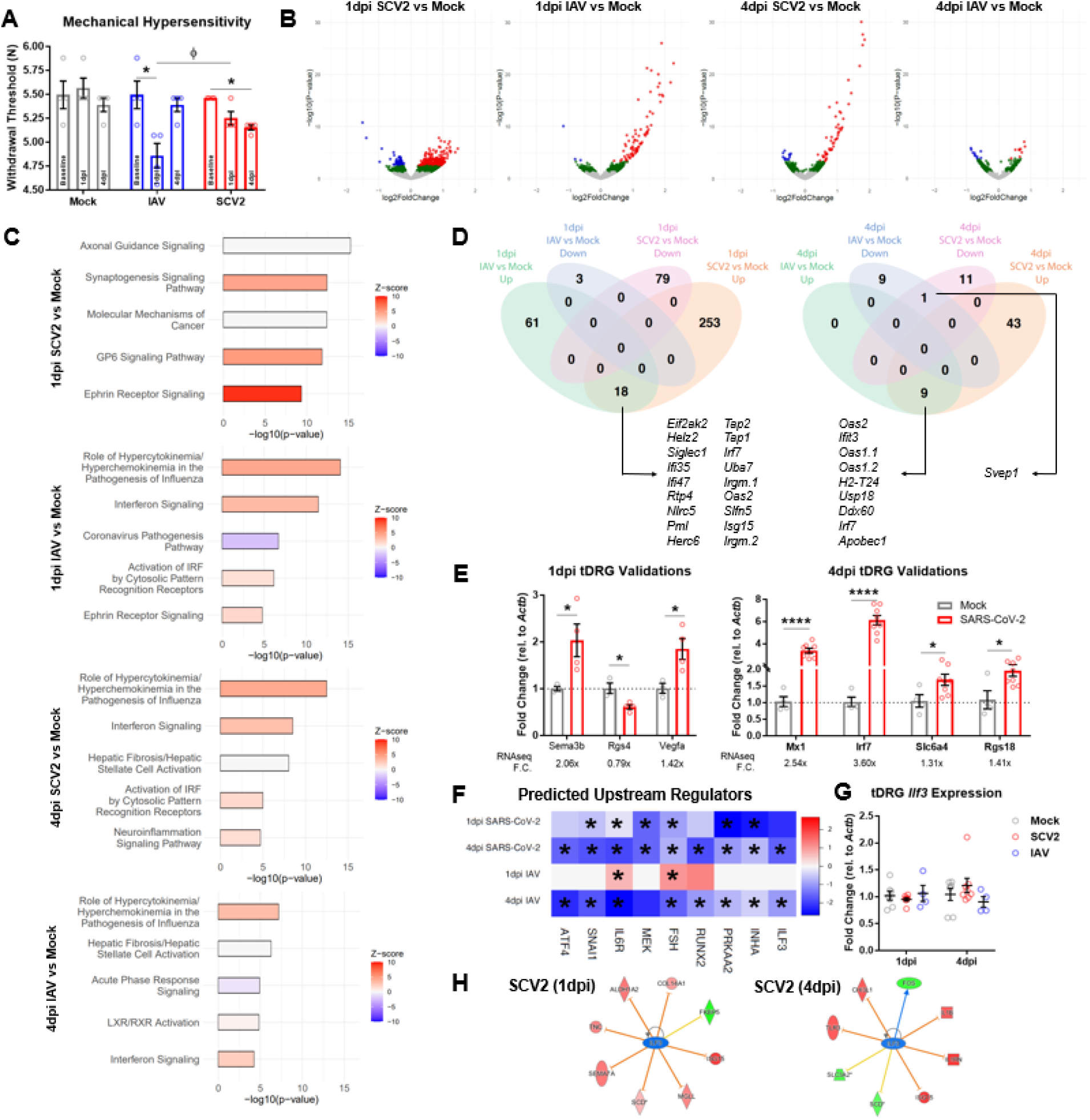
SARS-CoV-2 induces a unique behavioral phenotype and molecular signature in DRG tissue. **A**) Mechanical thresholds of mock, IAV, and SARS-CoV-2 animals at baseline, 1dpi, and 4dpi. IAV induced severe hypersensitivity at 1dpi, and SARS-CoV-2 induced mild hypersensitivity by 4dpi (n=4 per group; *p<0.05 for one-way ANOVA Tukey’s m.c.). IAV infection resulted in significantly lower thresholds than SARS-CoV-2 on 1dpi (Φp<0.05 for two-way ANOVA Tukey’s m.c.). **B**) Volcano plots for 1 and 4dpi SARS-CoV-2 and IAV tDRG RNA-seq (n=4 per group). Red=p-adj.<0.1, log2FC>0. Blue=p-adj.<0.1, log2FC>0. Green=p-nom.<0.05. **C**) Top 5 IPA Canonical Pathways for SARS-CoV-2 and IAV tDRGs (p-nom.<0.05; - log_10_(p-value)>1.3). **D**) Petal diagrams for 1dpi and 4dpi SARS-CoV-2 and IAV tDRG DEGs (p-adj.<0.01), with commonly upregulated or downregulated genes. **E**) qPCR validation of 1dpi and 4dpi SARS-CoV-2 tDRG DEGs (n=3-8 per group; *p<0.05, ****p<0.0001 for multiple t-tests). **F**) IPA predicted upstream regulators that had predicted inhibition in 1 and 4dpi SARS-CoV-2 tissues and 4dpi IAV tissues (*Benjamini-Hochberg p<0.05). **G**) No significant changes in *Ilf3* gene expression were observed in SARS-CoV-2, IAV, or Mock tDRGs 1 and 4dpi in accordance with sequencing (n=4-8 per group). **H**) IPA prediction of ILF3-regulated genes at 1 and 4dpi in SARS-CoV-2 tDRGs.

### Sensory Transcriptional Response to SARS-CoV-2 Infection

We conducted transcriptional profiling via RNA-seq on thoracic DRGs from SARS-CoV-2- and IAV-infected hamsters at both 1dpi and 4dpi because of their respiratory, visceral, and dermal innervations. Differential expression analysis of RNA-seq data revealed transcriptomic changes in both SARS-CoV-2- and IAV-infected thoracic DRGs compared to mock at 1dpi and 4dpi. SARS-CoV-2 infection resulted in a more robust differential expression at both time points: 344 genes at 1dpi (271 up & 79 down; p-adj.<0.1) and 63 genes at 4dpi (52 up & 11 down; p-adj.<0.1). IAV infection resulted in differential expression of 82 genes at 1dpi (79 up & 3 down; p-adj.<0.1) and 18 genes at 4dpi (9 up & 9 down; p-adj.<0.1) (**Figure 3B**). Considering the milder acute mechano-sensitivity phenotype in SARS-CoV-2-infected hamsters and greater differential gene expression compared to IAV-infected hamsters, we hypothesized that certain acute SARS-CoV-2-induced transcriptional changes may counteract interferon-induced somatosensory sensitization, potentially by causing a stronger neuronal gene adaptation signature. To better assess this, we performed a canonical pathway analysis (IPA, Qiagen) on our RNA-seq data. This analysis showed neuron-specific transcriptional differences within the reported top upregulated canonical pathways (based on genes with nominal p<0.05) (**Figure 3C**). The top two most enriched pathways for 1dpi SARS-CoV-2 tissue was “Axonal Guidance Signaling” and “Synaptogenesis Signaling”, and at 4dpi “Neuroinflammation Signaling” was among the top-five pathways. However, for IAV samples, the top canonical pathway results were consistently representative of generic viral response pathways.

To better understand which transcripts were driving these enriched annotations, we compared DEGs (p-adj.<0.1) between tissues derived from IAV- and SARS-CoV-2-infected hamsters. Commonly upregulated genes between 1dpi and 4dpi SARS-CoV-2 and IAV tissues were primarily anti-viral in nature, with only one co-downregulated gene emerging at 4dpi, *Svep1* (a vascular gene whose locus has been associated with poor SARS-CoV-2 clinical outcomes (*54*)) (**Figure 3D**). RNA-seq was validated at 1dpi and 4dpi through qPCR measurement of neuronal and anti-viral genes from SARS-CoV-2 and mock tissues. Interestingly, we observed bi-directional regulation of neuropathy-associated and/or pro-nociceptive genes at 1dpi, such as upregulation of *Sema3b (55) and Vegfa* (*56, 57*) and downregulation of *Rgs4* (*58*) (**Figure 3E**). qPCR validations of 4dpi included upregulation of *Mx1* and *Irf7* (pro-inflammatory, anti-viral genes) (*59*), as well as *Slc6a4* (*60*) and *Rgs18* (*61*),which have also been implicated in sensory abnormalities (**Figure 3E**).

Analysis of upstream regulators (URs; IPA, Qiagen) of differentially expressed nominal p<0.05 genes on IPA revealed several commonly- and oppositely-regulated URs between SARS-CoV-2 and IAV datasets. Based on our hypothesis that SARS-CoV-2 transcriptionally counteracts interferon-induced hypersensitivity, we wanted to identify URs uniformly associated with timepoints of acute viral infection during which lower levels of hypersensitivity were observed, namely 1dpi SARS-CoV-2, 4dpi SARS-CoV-2, and 4dpi IAV. We focused on URs with predicted downregulated activity in an attempt to find inhibition targets. Nine URs met this criterion: Interleukin 6 Receptor (IL6R), Mitogen-activated Protein Kinase Kinase (MEK), Interleukin Enhancer-binding Factor 3 (ILF3), Runt-related Transcription Factor 2 (RUNX2), Protein Kinase AMP-Activated Catalytic Subunit Alpha 2 (PRKAA2) (UR was AMPKα2 gene), Follicle Stimulating Hormone (FSH), Activating Transcription Factor 4 (ATF4), Snail Family Transcriptional Repressor 1 (SNAI1), and Inhibin Subunit Alpha (INHA) (**Figure 3F**). Interestingly, pre-clinical and clinical literature supports a positive association between upregulation/activation of IL6R (*62–64*), MEK (*65–67*), RUNX2 (*68, 69*), FSH (*70*), & ATF4 (*71, 72*) and nociceptive states, and several laboratories have validated interventions in relevant pathways as promising anti-nociceptive therapeutic strategies. Only AMPKα2 activity was expressed towards a pro-nociceptive direction in this list, as pre-clinical literature suggests activation of this protein is associated with the alleviation of nociceptive symptoms (*73, 74*).These data suggest that other targets in this list may serve as novel therapeutic avenues of pain management. Among the identified genes that have not been studied in pain models (SNAI1, ILF3, and INHA), we selected to study ILF3 as there is a commercially available inhibitor, YM155, which can be systemically applied and has been clinically tested in various cancer subtype populations (*75–77*).

Predicted interactions between ILF3 and SARS-CoV-2-regulated genes further support its investigation as a pain target, as several genes were associated with either neuronal activity/plasticity (including *Fos, Col14a1, Aldh1a2, Fkbp5, Sema7a, Mgll Chi3l1, and Slc3a2*),or with interferon and cytokine responses (including *Isg15, Il1b, Il1rn, Tlr3, Tnc*) (**Figure 3H**). As expected based on RNAseq, which did not label *Ilf3* as a significant DEG, whole tissue qPCR demonstrated a lack of *Ilf3* gene expression changes in SARS-CoV-2 and IAV tissues from 1dpi or 4dpi timepoints, suggesting that changes in the activity of this molecule are occurring at the protein level (**Figure 3G**). Of note, YM155 is believed to affect subcellular localization of ILF3 and its associated complexes, as opposed to directly inhibiting its expression (*78*).

### Inhibition of ILF3 Activity Alleviates Sensory Hypersensitivity in an Inflammatory Pain Model

We next used the CFA model of peripheral inflammation in female mice in order to determine the impact of ILF3 inhibition in sensory hypersensitivity behaviors associated with inflammatory pain states. We observed lethal toxicity at 20 mg/kg, so we proceeded with a 5 mg/kg once-daily regimen. In order to identify any immediate analgesic effects of YM155 under local, peripheral inflammation conditions, we first tested CFA-injected mice in the Von Frey and Hargreave’s assays at 30 minutes post-drug administration. YM155-treated mice displayed increased Hargreave’s response times (**Figure 4A**; RM two-way ANOVA Interaction: F(2,20)=4.116, df=2, p=0.0318; Sidak’s m.c.: YM155 vs Saline D3 Post-CFA (+drug) t=3.085, df=30, p=0.013; YM155 D2 Post-CFA (-drug) vs D3 Post-CFA (+drug) t=3.36, df=20, p=0.0186) and increased Von Frey thresholds (**Figure 4B**; RM two-way ANOVA Interaction: F(2,20)=13.5, df=2, p=0.0002; Sidak’s m.c.: YM155 vs Saline Day 4 Post-CFA (+drug) t=6.784, df=30, p<0.0001, YM155 D2 Post-CFA (-drug) vs D4 Post-CFA (+drug) t=8.517, df=20, p<0.0001). We also tested whether YM155 had sustained effects on sensory hypersensitivity after the expected window of activity (approximately 24 hours post-injection, based on a ~one hour half-life in intravenously-treated mice(*79*)). Indeed, when mice were monitored in the Hargreave’s assay at 24 hours post-injection, we observed a significantly higher withdrawal latency at six consecutive days (PD-D6) of YM155 administration (**Figure 4C**; RM two-way ANOVA Interaction: F(4,40)=2.887, df=4, p=0.0343; Sidak’s m.c.: YM155 vs Saline PD-D6 t=3.964, df=50, p=0.0012), prior to the expected recovery from thermal hypersensitivity in CFA animals. Similarly, we observed sustained recovery of mechanical thresholds on PD-D5, PD-D7, and PD-D9 in the Von Frey assay (**Figure 4D**; RM two-way ANOVA Interaction: F(4,40)=2.171, df=4, p=0.0897; Sidak’s m.c.: YM155 vs Saline PD-D5 t=3.59, df=50, p=0.0038; PD-D7 t=3.058, df=50, p=0.0177; PD-D9 t=4.122, df=50, p=0.0007). We observed no changes in weight due to YM155 administration over the first 9 days of treatment (**Figure 4E**).

**Figure 4.**
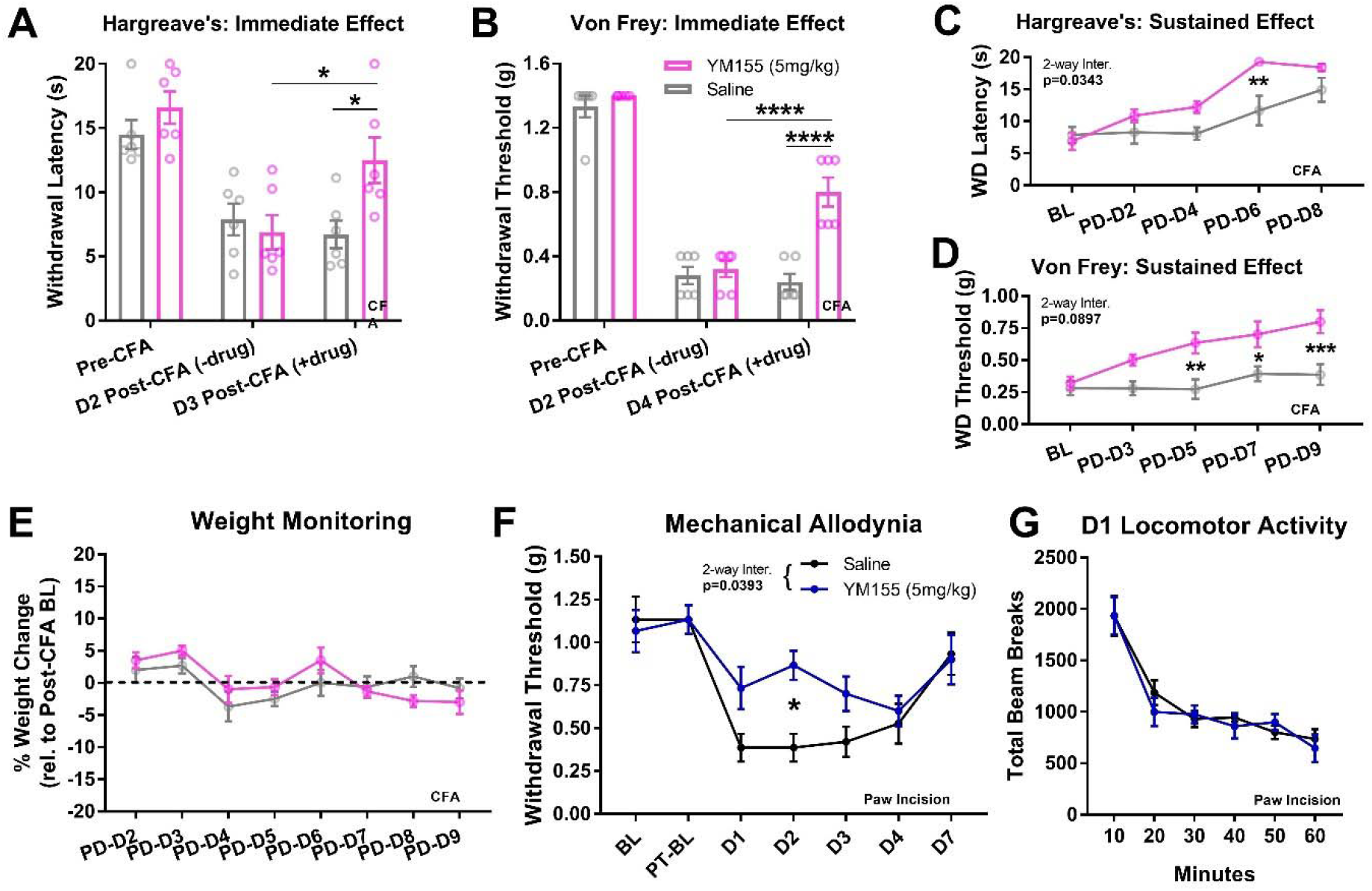
YM155 alleviates CFA-induced thermal and mechanical hypersensitivity in an immediate and sustained fashion. **A-B**) YM155 (5mg/kg i.p. QD) increased thermal and mechanical thresholds 30-60 minutes after administration (*p<0.05, ****p<0.0001 for two-way ANOVA Sidak’s m.c.) (n=6 per group). **C-D**) YM155 increased thermal and mechanical thresholds in a sustained fashion ~24 hours after administration by 5-6 days after initial administration (*p<0.05, **p<0.01, ***p<0.001 for two-way ANOVA Sidak’s m.c.) (n=6 per group). **E**) No changes in post-CFA weight were observed in YM155 animals throughout the course of administration (n=6 per group). **F**) Pre-treatment with YM155 led to significantly lower mechanical hypersensitivity after paw incision (*p<0.05 for two-way ANOVA Sidak’s m.c.) (n=6 per group). **G**) No differences in locomotion were observed on Day 1 post-paw incision between YM155 and Saline mice (n=6 per group).

We also tested whether YM155 could be used to prophylactically reduce pain experienced after acute post-operative injuries. For this, we used the paw incision model and pre-treated animals at a dose of 5mg/kg i.p. for seven days. Animals were not treated with drug after the incision. We observed a significant reduction in mechanical hypersensitivity due to the incision (**Figure 4F**; RM two-way ANOVA Interaction: F(6,60)=2.384, df=6, p=0.0393; Sidak’s m.c. YM155 vs Saline t=3.203, df=70, D2 p=0.0142). Importantly, we observed no changes in locomotor activity between animals immediately after testing mechanical hypersensitivity on D1 post-op (**Figure 4G**).

### SARS-CoV-2 Induces a Unique, Persistent Transcriptomic Profile in DRGs

Given that the severity of sensory hypersensitivity during acute infection with SARS-CoV-2 worsens over time and the existence of persistent sensory symptoms in patients afflicted by long COVID, we set out to determine whether the hamster respiratory model of SARS-CoV-2 infection displayed any prolonged sensory phenotypes. In this set of studies, we monitored mechanical hypersensitivity in male and female SARS-CoV-2, IAV, and mock treated hamsters at 28dpi (well-after viral clearance). Our findings reveal substantial mechanical hypersensitivity in SARS-CoV-2-infected hamsters of both sexes, but normal responses for IAV and mock hamsters (**Figure 5A**; for female groups: one-way ANOVA F(2,15)=8.469, p=0.0035, Tukey’s m.c. SARS-CoV-2vsMock q=5.385, df=15, p=0.0046; SARS-CoV-2vsIAV q=4.605, df=15, p=0.0138; for male groups: one-way ANOVA F(2,15)=22.36, p<0.0001, Tukey’s m.c. SARS-CoV-2vsMock q=8.043, df=15, p=0.0001; SARS-CoV-2vsIAV q=8.331, df=15, p<0.0001).

**Figure 5.**
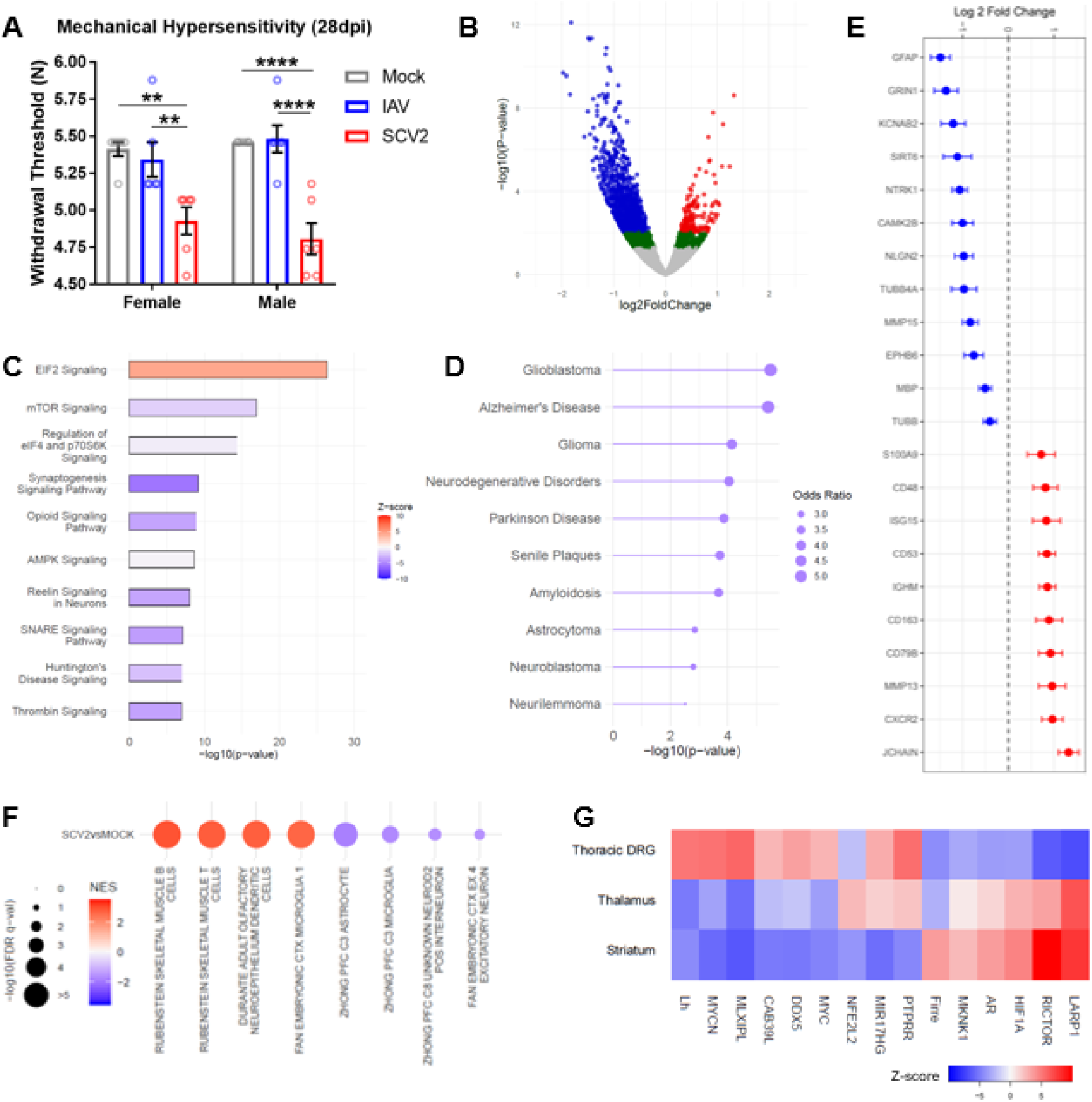
SARS-CoV-2 infection results in substantially reduced mechanical thresholds and a neuropathic transcriptomic landscape in the tDRGs well after viral clearance. **A**) Mechanical thresholds of mock, IAV, and SARS-CoV-2 animals at 28dpi (n=6 per group; **p<0.01, ****p<0.0001 for one-way ANOVA Tukey’s m.c.). **B**) Volcano plot for 31dpi SARS-CoV-2 tDRG RNA-seq (n=3 per group). Red=p-adj.<0.1, log2FC>0. Blue=p-adj.<0.1, log2FC>0. Green=p-nom.<0.05. **C**) IPA top 10 canonical pathways (-log10(p-value)>1.3) associated with 31dpi SARS-CoV-2 tDRG DEGs (p-nom.<0.05). **D**) enrichr DisGENET gateway top 10 diseases associated with 31 dpi SARS-CoV-2 tDRG DEGs (p-nom.<0.05). **E**) Log2(FC) of select neuronal and inflammatory genes from 31dpi RNA-seq (p-adj.<0.1). **F**) Positively and negatively enriched cell subtypes associated with 31dpi SARS-CoV-2 tDRG DEGs (GSEA NES>|1.5|; DEG p-adj.<0.1)**. G**) IPA top 15 upstream regulators between 31dpi SARS-CoV-2 tDRG, Striatum, and Thalamus (DEG p-nom.<0.05).

In order to determine whether longitudinally-altered DRG molecular mechanisms may be responsible for this SARS-CoV-2-specific hypersensitivity phenotype, we performed RNA-seq analysis and compared 31dpi thoracic DRGs between SARS-CoV-2 and Mock male animals. To our surprise, we identified 1065 DEGs (p-adj.<0.1, 170 up, 895 down; **Figure 5B**), which is a much larger number of DEGs than we observed with the 4dpi SARS-CoV-2 DRGs. Ontology analysis of DEGs (nominal p<0.05) also highlighted new and counter-regulated canonical pathways compared to those observed in 1dpi and 4dpi SARS-CoV-2 and IAV, including decreased “Synaptogenesis Signaling”, and the involvement of “EIF2 Signaling”, “mTOR Signaling”, “Opioid Signaling”, and “SNARE Signaling” (**Figure 5C**; −log_10_(p-value)>1.3). Furthermore, use of Enrichr’s DisGeNET gateway primarily associated these DEGs with neuro-oncological and neurodegenerative conditions, including Glioblastoma, Alzheimer’s Disease, Parkinson Disease, and Neurilemmoma (**Figure 5D**). Key DEGs (p-adj.<0.1) from RNAseq support our observed maladaptive alterations in canonical neuronal and inflammatory pathways, including changes in gene expression of several tubulin mRNA (*Tubb*) isoforms, myelin proteins, activity-related channels, extracellular matrix proteins, and cytokine/interferon-related proteins (**Figure 5E**).

Analysis of predicted cell subtype implications influence on 31dpi SARS-CoV-2 tDRG transcriptomic signatures using GSEA (C8 cell type signature gene set (v7.4)) revealed a positive contribution of pro-inflammatory cells, such as B cells, T cells, and dendritic cells (**Figure 5F**). Astrocytes, microglia, interneurons, and excitatory neurons contributions were negatively enriched (**Figure 5F**). Overall, these predictions suggest that 31dpi SARS-CoV-2 tDRGs are undergoing a pro-inflammatory state with inhibited neuronal and glial function, which is reflective of the ontology analysis above.

We next sought to determine whether a core group of upstream regulators (URs) may serve as a common target for sensory and perceptive components of pain, as well as affective comorbidities observed in long COVID-19 patients. We performed an IPA UR comparison analysis between our 31dpi DRG, Striatum, and Thalamus RNA-seq data, the latter two datasets coming from another systemic long-COVID study our group performed in hamsters under the same conditions (*32*). The Striatum and Thalamus are all well-cited regions involved in the initiation and maintenance of sensory components of pain, as well as emotional pain signs, such as catastrophizing (*80, 81*). Here, we focused on the top common upstream regulators across these regions.

Interestingly, a majority of the top 15 URs demonstrated a unidirectional predicted activation/inhibition state between Thalamus and Striatum, but not DRGs (**Figure 5G**). However, we did observe a common upregulation of PTPRR and miR17hg, as well as a downregulation of FIRRE, between DRG and Thalamus. While PTPRR, a protein tyrosine phosphatase receptor, has not been implicated in pain, human studies have suggested an association between its upregulation and depression (*82, 83*). *MIR17HG* (a long non-coding RNA (lncRNA) involved in cell survival) gene abnormalities have also been reported in Feingold 2 syndrome patients that suffer from chronic myofascial pain and affective symptoms (*84, 85*). FIRRE, another lncRNA, has been implicated in spinal cord neuropathic pain mechanisms (*86*).

Thus, common regulators between the peripheral and central nervous systems may serve as useful targets for both sensory and affective symptoms of long COVID-19.

### SARS-CoV-2 Infection Causes Transcriptomic Signatures Similar to Persistent Inflammation and Nerve Injury Models in Dorsal Root Ganglia

While our bioinformatic analysis of SARS-CoV-2 RNA-seq datasets led to the identification of potential treatment targets, such as ILF3, we also wanted to determine if a meta-analysis of this data against existing injury datasets may yield a more comprehensive list of pain targets. We therefore compared 1, 4, and 31dpi thoracic DRG RNA-seq from SARS-CoV-2-infected hamsters against GEO RNA sequencing data from the aforementioned murine SNI and CFA datasets.

We observed several commonly upregulated genes between SARS-CoV-2 and CFA at both 1 and 4dpi, and only on 1dpi when comparing to SNI (**Figure 6A**). Interestingly, we identified a group of 53 genes that were upregulated by SARS-CoV-2 at 1dpi but downregulated by SNI (**Figure 6A**). g:Profiler associated this gene set with neuroplasticity, particularly in the synaptic/dendritic cellular compartments, and strongly associated the Sp1 transcription factor (implicated in several pro-nociceptive mechanisms) with these genes (**Figure 6B**) (*87–89*). Some of these genes, such as *Scn4b (90, 91), Rhobtb2 (92), Mgll* (*93, 94*), and *Cntfr* (*95*) have been positively associated with sensory hypersensitivity under injury states, suggesting they may be unique mechanisms by which SARS-CoV-2 induces mild hypersensitivity. This finding also highlights potential SNI-induced compensatory anti-nociceptive gene programs. However, anti-nociceptive genes were also upregulated by SARS-CoV-2, including *Gprc5b* (*96*) and *Grk2* (*97, 98*). Several genes implicated in neurodevelopment and dendritic plasticity were also upregulated by SARS-CoV-2, but have not yet been studied in pain. Interesting candidates include *Olfm1, Fxr2, Atcay, Cplx1, Iqsec1, Dnm1, Clstn1, Rph3a, Scrt1, Ntng2* and *Lhfpl4*. Ontologies significantly associated with this SARS-CoV-2 versus SNI contra-regulated gene list are GO:BP nervous system development (p-adj=0.005994), GO:BP generation of neurons (p-adj=0.024), GO:CC somatodendritic compartment (p-adj=0.004204), GO:CC synapse (p-adj=0.01054), and GO:CC cell junction (p-adj=0.02067). We also identified a core set of genes, mostly associated with extracellular matrix remodeling, was commonly upregulated between 1dpi SARS-CoV-2, CFA, and SNI: *Col1a1, Col1a2, Col6a3, Hspg2, Irgm, Lama2, Lamb1, Lamc1, and Siglec1* (**Figure 6C**). This is in agreement with previous literature implicating extracellular matrix remodeling with the maintenance of inflammatory- and nerve injury-associated pain sensation (*99*).

**Figure 6.**
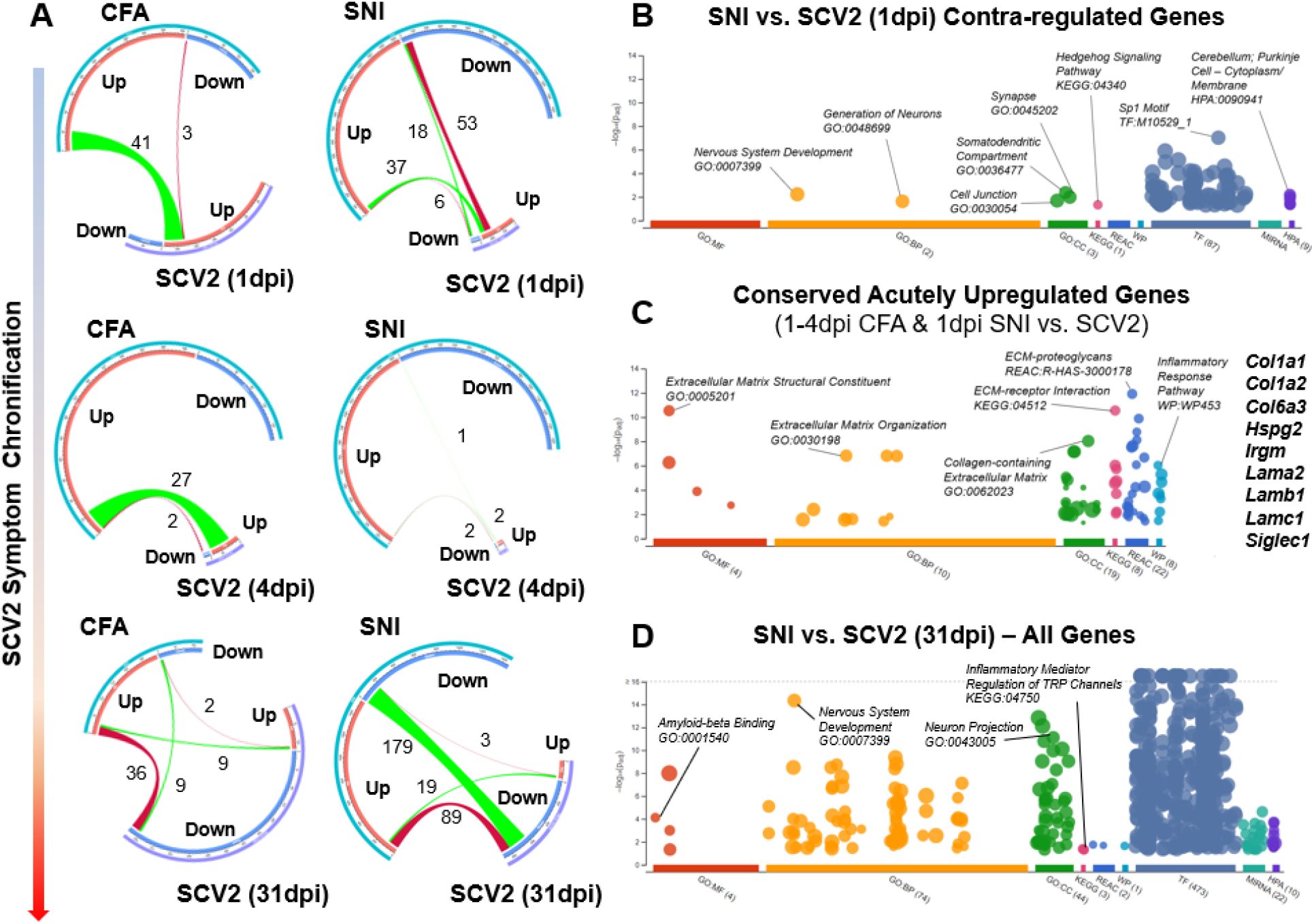
SARS-CoV-2 infection causes a longitudinally variable tDRG transcriptomic profile that shares pro-nociceptive components yet demonstrates unique plasticity signatures. **A**) Chord diagrams demonstrating regulation of DRG gene expression changes between CFA, SNI, and SARS-CoV-2 (1, 4, and 31dpi) animals. **B-D**) Dot plots demonstrating significant Gene Ontology (GO; Molecular Function, Biological Process, Cellular Compartment), Kyoto Encyclopedia of Genes and Genomces (KEGG), Reactome (REAC), WikiPathways (WP), Transfac (TF), and Human Protein Atlas (HPA) (-log10p-adj.>1.3) for contra-regulated genes between SNI and 1dpi SARS-CoV-2, conserved upregulated genes for CFA/SNI vs 1-4dpi SARS-CoV-2 comparisons, and all commonly regulated genes between SNI & 31dpi SARS-CoV-2. Dot plots adapted from g:Profiler.

Lastly, comparison of all genes regulated by CFA and 31dpi SARS-CoV-2 revealed a subset of counter-regulated DEGs (36 CFA Up-SARS-CoV-2 down; p-adj.<0.1). These genes are implicated in pathways such as myelination/axon ensheathment (*Mpz, Mbp, Prx, Fa2h, Dhh*, and *Mag*), semaphorin-regulation of axonogenesis (*Sema3g and Sema4g*), and extracellular matrix organization (*Nid2, Col5a3, Mmp15, Mmp14, Col4a1, and Fscn1*) (g:profiler GO:BP p-adj.<0.05). We observed a strong transcriptional counter-regulation between SNI and 31dpi SARS-CoV-2 as well (89 SNI up-SARS-CoV-2 down; p-adj.<0.1). This signature was predominantly related with nervous system development, with implicated genes including *Mpz, Plec, Prkcg Metrn, Slit1, Brd2, Anks1a, Cpne5, Sema4f, Hspg2, Sh3gl1, Prag1, Map6, Mdga1, Fphs, Ppp2r5b, Plod3, Phgdh, Dpysl5, Gpc1, Elavl3, Gpsm1, Marcksl1, Col4a1, Niban2, Carm1, Irs2, Lgi4, Erbb2, Syngap1*, and *Nlgn2* (g:profiler GO:BP p-adj.<0.05).

However, we were mostly surprised by the robust overlap of downregulated DEGs between SNI and 31dpi SARS-CoV-2 (179; p-adj.<0.1). Nervous system development and morphogenesis were robust pathway signatures, implicating neuronal plasticity as a key contributor to nerve injury and virus-induced pain states. But this comparison also uniquely revealed strongly altered synaptic transmission pathways, with DEGs including *Slc7a7, Syngr1, Prkaca, Rab3a, Ntrk1, Nptx1, Stx1b, Jph3, Mapk8ip2, Calm3, Pnkd, Ppp1r9b, Pip5k1c, Cacng7, Dlgap3, Nrxn2, Pink1, Grk2, Ncdn, Cplx2, Camk2b, Grin1, Brsk1, Ache*, and *Jph4*. This gene list suggests that SARS-CoV-2 mirrors nerve injury maladaptive mechanisms both through direct modification of neuronal excitability at the membrane level and through modulation of transcriptional regulation elements. These, along with other implicated pathways from the overall SNI-SARS-CoV-2 31dpi comparison, such as amyloid-beta binding and TRP channel modulation, are highlighted in **Figure 6D**.

Combined, this meta-analysis emphasizes SARS-CoV-2’s ability to recapitulate transcriptional perturbations in the DRG underlying both inflammatory- and nerve injury-associated pain states. However, these findings also demonstrate the induction of plasticity-associated perturbations that counter those seen in other injury models. Future studies will elucidate whether these differences promote the maintenance of mechanical hypersensitivity we observed in SARS-CoV-2-infected animals. Furthermore, these findings support the use of the SARS-CoV-2 respiratory infection hamster model as a preclinical chronic pain model, which can be used for the understanding of the evaluation of pharmacological treatments.

## DISCUSSION

The relatively high prevalence of both acute asymptomatic SARS-CoV-2 cases and positive somatosensory abnormalities in long COVID-19 patients prompted our group to investigate the ability of SARS-CoV-2 to perturb sensory nervous system functions. By utilizing the established golden hamster model of COVID-19 (*100, 101*), we detected low levels of SARS-CoV-2-derived RNA in the absence of infectious particles. Exposure of sensory tissues to this viral material and/or the resulting type I interferon response correlated with a progressive and prolonged mechanical hypersensitivity signature that was unique to SARS-CoV-2. Transcriptomic analysis of thoracic SARS-CoV-2-infected DRGs highlighted a pronounced neuronal signature unlike the predominantly pro-inflammatory signature seen in IAV-infected DRGs. SARS-CoV-2 infection also correlated with worsened hypersensitivity post-recovery in both female and male hamsters, which may be attributable to altered excitability, cytoskeletal architecture, extracellular remodeling, and myelination as a result of the host response to this inflammatory material. Transcriptional profiling of tDRGs at 1, 4, and 31dpi implicated several potential therapeutic targets for the management of chronic pain. Indeed, the prediction of ILF3 inhibition as a potential therapeutic intervention was validated in the murine CFA model of peripheral inflammation. Lastly, meta-analysis against existing transcriptional data sets from pain models highlighted several unexplored acutely and chronically contra-regulated genes between SARS-CoV-2 and SNI that could serve as future targets for anti-nociceptive therapies and provide novel mechanistic insight into these perturbations.

The SARS-CoV-2 RNA infiltration dynamics within sensory tissue observed in this study were similar to those noted in our longitudinal study of SARS-CoV-2 effects on the brain, where a rapid transcriptional induction to infection is followed by a return to baseline in most, but not all tissues (*32*). While we confirmed the presence of SARS-CoV-2 RNA in various cell types of the DRG, we were surprised by the neuronally-biased transcriptional responses associated with this positivity that were not as prominent in tissue from IAV-infected hamsters. Together, these data suggest that the host response to SARS-CoV-2 infection elicits a unique transcriptional output capable of inducing lasting changes to DRG plasticity.

In addition to elucidating the impact SARS-CoV-2 has on DRGs, this study also identified a subset of host factors as modulators of the nociceptive responses. Of note, increased activity of ILF3 (*102–104*) is generally considered oncogenic. Furthermore, several of the disease risk signatures associated with gene changes observed in the 31dpi SARS-CoV-2 DRGs revolved around neuronal and glial cancers. Given our group and other’s current (ILF3 inhibitor) and prior (*Rgs4* downregulation (*80*), HDAC1 inhibition (*105*), and HDAC6 (*106*) inhibition) successes with use of cancer-targeting therapies for the treatment of inflammatory- and nerve injury-associated pain states, we believe that the careful repurposing of existing clinically-validated cancer therapeutics may serve as one possible strategy for providing alternative treatments for pain management. Implementation of this treatment strategy will necessitate molecular modifications or intricate drug delivery strategies to reduce potential systemic toxicities.

Future studies will focus on robust characterization of predicted pathways and validation of novel treatment interventions. For example, Ephrin Receptor Signaling, which frequently appeared in our ontology analyses, has a documented role in nociceptive processing (*107*). Ephrin signaling is an essential mediator of extracellular matrix dynamics (*108*), which subsequently affect synaptic plasticity in the form of neurite outgrowth and synaptic integrity (*109*). Current pain therapeutics are primarily focused on modulating maladaptive neuronal hyperexcitability through GPCR or ion channel targeting (*110*). However, few interventions target downstream transcriptomic mechanisms that broadly influence synaptic plasticity, an essential component of central sensitization. Along with ILF3, upstream regulators of the SARS-CoV-2-activated Ephrin pathway may support this alternative treatment direction.

Furthermore, few pain therapeutics target both the peripheral and central site of the nociceptive pathway. Here, we identified that several common predicted upstream regulator targets exist between the DRGs and brain regions that process pain and emotion. While most of the top common URs were counter-regulated between the DRGs and Thalamus/Striatum, three promising pain- and affect-associated URs (PTPRR, miR17HG, and FIRRE) were predicted to change unidirectionally between DRG and Thalamus. Work from our group has shown a high level of treatment effectiveness in targeting the same protein in DRG and Thalamus through studies on the signal transduction modulator RGS4 (*80 and unpublished*). Notably, in this study, the expression of the *Rgs4* gene was decreased in 1dpi in DRGs of SARS-CoV-2-infected hamsters.

Finally, while several groups have recapitulated human COVID-19 symptoms in this respiratory hamster model, this is the first study that confirmed the model’s relevance for somatosensory symptoms. From a mechanical hypersensitivity perspective, we believe this model accurately aligns with the somatosensory trajectory of many COVID-19 patients, both acutely and chronically. This SARS-CoV-2 model was also useful for further identifying core mechanisms across pain models, while also potentially providing insights into novel viral-mediated nociceptive states with relevance for drug development.

## METHODS

### Infection & Local Inflammation Animal Models

One- to two-month-old male golden hamsters (*Mesocricetus auratus*) were used in all infection experiments, and age-matched female hamsters were included in 31dpi experiments (Charles River Laboratories, MA). Male hamsters were co-housed on a twelve-hour light-dark cycle and had access to food and water *ad libitum*. Female hamsters were housed individually to prevent injury due to aggression. Hamster work was performed in a CDC/USDA-approved biosafety level 3 laboratory in accordance with NYU Langone and Icahn School of Medicine at Mount Sinai IACUC protocols. Mice were housed on a twelve-hour light-dark cycle and had access to food and water *ad libitum* in accordance with the Icahn School of Medicine at Mount Sinai IACUC protocols.

Two- to three-month-old hamsters received an intranasal inoculation of 100μL of phosphate-buffered saline (PBS) containing 1000 plaque forming units (PFU) of SARS-CoV-2, 100,000 PFU of IAV (viral control), or PBS alone (mock control). Hamsters were euthanized by intraperitoneal pentobarbital injection followed by cardiac perfusion with 60 mL PBS.

For studies using models of peripheral inflammation, two-to three-month old mice received 30uL left hindpaw injections of Complete Freund’s Adjuvant (CFA; diluted 1:1 in saline), as described (*106*). For studies using the post-operative incision model, two- to three-month old mice received an incision from the posterior plantar surface of the hindpaw to the middle of the paw pads, in which dermis and superficial muscle was cut and dermis was sutured afterwards as cited (*111*). CFA and paw incision groups of mice received daily intraperitoneal (i.p.) injections of saline (vehicle) or YM155 (Tocris Biosciences), an Interleukin Enhancer Binding Factor 3 (ILF3) inhibitor (5mg/kg diluted in saline).

### Von Frey Assay

Hamsters/mice were placed on a raised grid platform in plastic containers and were allowed to habituate to their environment for a minimum of 10 minutes. Afterwards, filaments of ascending forces were applied to the left hindpaw and responses were recorded. A positive response consisted of a hindpaw lift, shake, or lick. Progression to the next filament was determined by recording of positive or negative responses for three out of five applications with each filament. Mechanical withdrawal threshold was defined as the first (for hamsters, to minimize cross-contamination of cohorts by prolonged fomite exposure) or second (mouse, for consistency) filament force at which an animal had three positive responses. All materials utilized for testing of infected hamsters were thoroughly decontaminated between testing of infection groups.

### Hargreave’s Assay

The CFA model induces thermal hypersensitivity for 10-14 days on average (*80*). We used the Hargreave’s thermal beam assay to assess the effects of YM155 administration on thermal hypersensitivity associated with left hindpaw CFA injection. Mice were placed on a Hargreave’s platform in plastic containers and were allowed to habituate for 30 minutes. A light beam heat source (IITC Life Science Inc., CA) set to an intensity level of IF=30 was aimed at the left hindpaw for a maximum of 20 seconds (cutoff). Similarly to Von Frey, paw withdrawal was defined as a hindpaw lift, shake, or lick. Three measurements were recorded and averaged for each hindpaw, with each measurement taking place at least two minutes apart.

### Tissues

Tissues were harvested at 1, 4, and 31 dpi and immediately placed in TRIzol (Invitrogen, MA) for transcriptomic analysis or 4% paraformaldehyde (PFA) in phosphate-buffered saline (PBS) for histology or fluorescent *in situ hybridization* (RNAscope). Fixed tissues were sucrose converted after 48 hours of 4% PFA fixation in 10% sucrose in PBS (Day 1), 20% sucrose in PBS (Day 2), and 30% sucrose in PBS with 0.01% azide (Day 3). Slide-mounted tissues were paraffin-embedded and sliced to a thickness of 5 microns. Tissue collected for transcriptomic analysis were homogenized in Lysing Matrix A homogenization tubs (MP Biomedicals, CA) for two cycles (40s; 6m/s) in a FastPrep 24 5g bead grinder and lysis system (MP Biomedicals, CA). Tissue collected for plaque assays was homogenized in 1 mL PBS in Lysing Matrix A homogenization tubs (MP Biomedicals, CA) for two cycles (40s; 6m/s).

### RNA Isolation & qPCR

RNA was isolated through a phenol:chloroform phase separation protocol as detailed in the TRIzol Reagent User Guide. RNA concentrations were measured by NanoDrop (Thermofisher, MA). 1,000ng of cDNA was synthesized using the qScript cDNA Synthesis kit (QuantaBio, MA) as detailed in the qScript cDNA Synthesis Kit Manual. Exon-exon-spanning primers targeting as many splice variants as possible were designed with Primer-BLAST (National Center for Biotechnology Information, MD). qPCRs were performed in triplicate with 30 ng of cDNA and a master mix of exon-spanning primers (Supplementary Table 1) and PerfeCTa SYBR Green FastMix ROX (QuantaBio, MA) on an QuantStudio real-time PCR analyzer (Invitrogen, MA), and results were expressed as fold change (2^-ΔΔCt^) relative to the β-actin gene (*Actb*).

### Plaque Formation Assay

Plaque assays were performed as described previously (*31*). Virus was logarithmically diluted in SARS-CoV-2 infection medium with a final volume of 200 uL volume per dilution. 12-well plates of Vero E6 cells were incubated for 1 hour at room temperature with gentle agitation every 10 minutes. An overlay comprised of Modified Eagle Medium (GIBCO), 4 mM L-glutamine (GIBCO), 0.2% BSA (MP Biomedicals), 10 mM HEPES (Fisher Scientific), 0.12% NaHCO3, and 0.7% Oxoid agar (Thermo Scientific) was pipetted into each well. Plates were incubated at 37 degrees C for 48 hours prior to fixation in 4% PFA in PBS for 24 hours. Plaques were visualized via staining with crystal violet solution (1% crystal violet (w/v) in 20% ethanol (v/v)) for 15 minutes.

### RNAscope In Situ Hybridization

The Fluorescent Multiplex V2 kit (Advanced Cell Diagnostics, CA) was used for RNAscope FISH. Specifically, we used the FFPE protocol as detailed in the RNAscope Multiplex Fluorescent Reagent Kit v2 Assay User Manual. RNAscope probes were as follows: *Rbfox3* (NeuN) for pan-neuronal labeling (Mau-Rbfox3-C1) and the Spike gene (*S*) for SARS-CoV-2 labeling (V-nCoV2019-S-C3). Opal dyes (Akoya Biosciences, MA) were used for secondary staining as follows: Opal 690 for C1 and Opal 570 for C3. DAPI was used for nuclear staining. Images were taken on an LSM880 confocal microscope (Zeiss, GER) with identical parameters between mock and SARS samples.

### Immunohistochemistry

Immunohistochemistry was performed according to protocols described previously (*32*).Briefly, 5μm sections were cut from FFPE tissues and mounted on charged glass slides. Sections were deparaffinized by immersion in xylene and subsequently submerged in decreasing concentrations of ethanol to rehydrate. Rehydrated sections were submerged in IHC-Tek Epitope Retrieval Solution (Cat #IW-1100) and steamed for 45min in IHC-Tek Epitope Retrieval Steamer (Cat #IW-1102) for antigen retrieval. Tissues were blocked with 10% goat serum and 1% bovine serum albumin in TBS for 1hr at room temperature. Primary antibody (monoclonal murine-derived anti-SARS-CoV-2 N protein) was diluted 1:100 in a 1% BSA TBS solution and added to slides. Slides were incubated with primary antibody solution overnight at 4°C. Slides were washed in TBS with 0.025% Triton-X-100 and treated with 0.3% hydrogen peroxide in TBS for 15min. Slides were washed once again. HRP-conjugated goat anti-mouse IgG secondary antibody (ThermoFisher, Cat #A21426) was diluted 1:5000 and added to slides. Slides incubated with secondary antibody at room temperature for 1hr. Slides were washed twice, and DAB developing reagent (Vector Laboratories, Cat #SK-4105) was added to slides. Slides were dehydrated with increasing concentrations of ethanol and cleared using xylene. Slides were cover slipped, dried, and imaged using brightfield setting on EVOS M5000 inverted microscope.

### RNA Sequencing

RNA was isolated from tissues as previously described above. 500ng-1μg of total RNA per sample was enriched for polyadenylated RNA and prepared for RNA sequencing using the TruSeq Stranded mRNA Library Prep Kit (Illumina) per manufacturer instructions. Samples were sequenced on an Illumina NextSeq 500 platform or by the NYU Langone Genome Technology Center. FASTQ files were then aligned to the golden hamster genome (MesAur 1.0, ensembl) via the RNA-Seq Alignment application (BaseSpace, Illumina). Salmon files were analyzed using DESeq2 (*112*). For non-ontology analyses, all genes with an adjusted p-value (p-adj) less than 0.1 were considered “Differentially Expressed Genes” (DEGs).

Ontological analysis was performed using g:Profiler and Qiagen Ingenuity Pathway Analysis, targeting genes with a nominal p-value of less than 0.05 to increase analytical power. All visualizations of RNA-seq, differential expression analysis, and ontological analysis data were created by the respective ontological analysis programs or by R using ggplot2, VennDiagram, Circos, pheatmap, ComplexHeatmap, and gplots packages.

Gene set enrichment analyses were conducted using the GSEA Java application for Mac (v 4.1.0) (MSigDB; Broad Institute, UC San Diego). Analyses were performed on pre-ranked gene lists derived from differential expression data. Genes were ranked by the following statistic: −log10(p-value)/sign(log2FoldChange). GSEA analyses were conducted against the C8 cell type signature gene set (v7.4) provided by the Molecular Signatures Database (MSigDB).

### Meta-Analysis

FASTQ files from Parisien et al. (2019) (*99*) generated from RNA-seq of DRG tissues from mice subjected to sham (mock), Complete Freund’s Adjuvant (CFA), and Spared Nerve Injury (SNI) treatments were obtained from NCBI GEO (GSE111216). Paired end read files were aligned to the *Mus musculus* transcriptome (GRCm39) and quantified using Salmon (version 1.4.0). Salmon files were analyzed for differentially expressed genes using DESeq2, and all genes expressing a p-adj<0.1 were considered differentially expressed. Differentially expressed genes from murine DRG injury models compared to mock tissues were compared to analogous differentially expressed genes from infected hamster DRG tissues compared to mock hamster DRG tissues. These comparative analyses were visualized using Circos, VennDiagram, and ggplot2. Shared and contra-regulated gene sets highlighted from these analyses were also analyzed for ontology using g:Profiler.

### Statistical Analyses

All statistical analyses outside of sequencing-related assays were performed in GraphPad Prism Version 10. Repeated measure one- and two-way ANOVAs were used to compare the effects of virus type and time of infection on mechanical hypersensitivity, and post-hoc Tukey’s multiple comparison test were used to perform timepoint comparisons for the Von Frey assay. Multiple t-tests and two-way ANOVAs were used for qPCR analysis.

RNA-seq data was analyzed as described above. Ontology analysis statistics were performed with either Ingenuity Pathway Analysis (IPA), g:Profiler, or Enrichr (*113*).

## ACKNOWLEDGEMENTS

We would like to further thank Francis Avila, Virginia Gillespie, DVM, Ying Dai, and the rest of the staff at the Mount Sinai Center for Comparative Medicine and Surgery and the Mount Sinai Biorepository and Pathology Core for their technical assistance in tissue preparation for histology.

## AUTHOR CONTRIBUTIONS

Study concept and design: RAS, JJF, BT, VZ. Tissue Harvesting: RAS, JJF, KDP. Quantitative PCR and Plaque Assays: RAS, JJF, I. Golynker. RNAscope & IHC: JZ & JJF. RNA-seq & Data Analysis: RAS, JJF. Behavioral Assays: RAS, JJF, I. Giosan. Supervision: BT, VZ. Drafting of original manuscript: RAS, JJF. All authors reviewed, revised, and approved the final version of this paper.

## DATA AND MATERIALS AVAILABILITY

RNAseq data will be uploaded to NCBI GEO upon prior to publication of the manuscript for public accessibility. Other data may be provided upon request.

## References

1. G. Pascarella, A. Strumia, C. Piliego, F. Bruno, R. Del Buono, F. Costa, S. Scarlata, F. E. Agrò, COVID-19 diagnosis and management: a comprehensive review J. Intern. Med. 288 (2020), doi:10.1111/joim.13091.

2. E. Caronna, A. Ballvé, A. Llauradó, V. J. Gallardo, D. María Ariton, S. Lallana, S. L. Maza, M. O. Gadea, L. Quibus, J. L. Restrepo, M. Rodrigo-Gisbert, A. Vilaseca, M. H. Gonzalez, M. M. Gallo, A. Alpuente, M. Torres-Ferrus, R. P. Borrell, J. Alvarez-Sabin, P. Pozo-Rosich, Headache: A striking prodromal and persistent symptom, predictive of COVID-19 clinical evolution. Cephalalgia 40 (2020), doi:10.1177/0333102420965157.

3. M. Amanat, N. Rezaei, M. Roozbeh, M. Shojaei, A. Tafakhori, A. Zoghi, I. A. Darazam, M. Salehi, E. Karimialavijeh, B. S. Lima, A. Garakani, A. Vaccaro, M. Ramezani, Neurological manifestations as the predictors of severity and mortality in hospitalized individuals with COVID-19: a multicenter prospective clinical study. BMC Neurol. 21 (2021), doi:10.1186/s12883-021-02152-5.

4. S. Escalard, V. Chalumeau, C. Escalard, H. Redjem, F. Delvoye, S. Hébert, S. Smajda, G. Ciccio, J. P. Desilles, M. Mazighi, R. Blanc, B. Maïer, M. Piotin, Early brain imaging shows increased severity of acute ischemic strokes with large vessel occlusion in COVID-19 patients. Stroke (2020), doi:10.1161/STROKEAHA.120.031011.

5. M. K. Halushka, R. S. Vander Heide, Myocarditis is rare in COVID-19 autopsies: cardiovascular findings across 277 postmortem examinations. Cardiovasc. Pathol. 50 (2021), doi:10.1016/j.carpath.2020.107300.

6. J. S. Stevens, K. L. King, S. Y. Robbins-Juarez, P. Khairallah, K. Toma, H. A. Verduzco, E. Daniel, D. Douglas, A. A. Moses, Y. Peleg, P. Starakiewicz, M. T. Li, D. W. Kim, K. Yu, L. Qian, V. H. Shah, M. R. O’Donnell, M. J. Cummings, J. Zucker, K. Natarajan, A. Perotte, D. Tsapepas, K. Krzysztof, G. Dube, E. Siddall, S. Shirazian, T. L. Nickolas, M. K. Rao, J. M. Barasch, A. M. Valeri, J. Radhakrishnan, A. G. Gharavi, S. A. Husain, S. Mohan, High rate of renal recovery in survivors of COVID-19 associated acute renal failure requiring renal replacement therapy. PLoS One 15 (2020), doi:10.1371/journal.pone.0244131.

7. X. J. Song, D. L. Xiong, Z. Y. Wang, D. Yang, L. Zhou, R. C. Li, Pain Management during the COVID-19 Pandemic in China: Lessons Learned Pain Med. (United States) (2020), doi:10.1093/PM/PNAA143.

8. M. A. Ellul, L. Benjamin, B. Singh, S. Lant, B. D. Michael, A. Easton, R. Kneen, S. Defres, J. Sejvar, T. Solomon, Neurological associations of COVID-19 Lancet Neurol. (2020), doi:10.1016/S1474-4422(20)30221-0.

9. S. Andalib, J. Biller, M. Di Napoli, N. Moghimi, L. D. McCullough, C. A. Rubinos, C. O’Hana Nobleza, M. R. Azarpazhooh, L. Catanese, I. Elicer, M. Jafari, F. Liberati, C. Camejo, M. Torbey, A. A. Divani, Peripheral Nervous System Manifestations Associated with COVID-19Curr. Neurol. Neurosci. Rep. (2021), doi:10.1007/s11910-021-01102-5.

10. A. Carfì, R. Bernabei, F. Landi, Persistent symptoms in patients after acute COVID-19 JAMA - J. Am. Med. Assoc. 324 (2020), doi:10.1001/jama.2020.12603.

11. K. Stavem, W. Ghanima, M. K. Olsen, H. M. Gilboe, G. Einvik, Persistent symptoms 1.5-6 months after COVID-19 in non-hospitalised subjects: A population-based cohort study. Thorax 76 (2021), doi:10.1136/thoraxjnl-2020-216377.

12. A clinical case definition of post COVID-19 condition by a Delphi consensus, 6 October 2021. World Heal. Organ. (2021).

13. C. H. Sudre, B. Murray, T. Varsavsky, M. S. Graham, R. S. Penfold, R. C. Bowyer, J. C. Pujol, K. Klaser, M. Antonelli, L. S. Canas, E. Molteni, M. Modat, M. Jorge Cardoso, A. May, S. Ganesh, R. Davies, L. H. Nguyen, D. A. Drew, C. M. Astley, A. D. Joshi, J. Merino, N. Tsereteli, T. Fall, M. F. Gomez, E. L. Duncan, C. Menni, F. M. K. Williams, P. W. Franks, A. T. Chan, J. Wolf, S. Ourselin, T. Spector, C. J. Steves, Attributes and predictors of long COVID. Nat. Med. (2021), doi:10.1038/s41591-021-01292-y.

14. H. E. Davis, G. S. Assaf, L. McCorkell, H. Wei, R. J. Low, Y. Re’em, S. Redfield, J. P. Austin, A. Akrami, Characterizing long COVID in an international cohort: 7 months of symptoms and their impact. EClinicalMedicine (2021), doi:10.1016/j.eclinm.2021.101019.

15. L. Sigfrid, T. M. Drake, E. Pauley, E. C. Jesudason, P. Olliaro, W. S. Lim, A. Gillesen, C. Berry, D. J. Lowe, J. McPeake, N. Lone, D. Munblit, A. Casey, P. Bannister, C. D. Russell, L. Goodwin, A. Ho, L. Turtle, M. E. O’Hara, C. Hastie, C. Donohue, R. G. Spencer, C. Donegan, A. Gummery, J. Harrison, H. E. Hardwick, C. E. Hastie, G. Carson, L. Merson, J. K. Baillie, P. Openshaw, E. M. Harrison, A. B. Docherty, M. G. Semple, J. T. Scott, Long Covid in adults discharged from UK hospitals after Covid-19: A prospective, multicentre cohort study using the ISARIC WHO Clinical Characterisation Protocol. Lancet Reg. Heal. - Eur. (2021), doi:10.1016/j.lanepe.2021.100186.

16. H. J. Zhao, X. X. Lu, Y. Bin Deng, Y. J. Tang, J. C. Lu, COVID-19: Asymptomatic carrier transmission is an underestimated problem. Epidemiol. Infect. (2020), doi:10.1017/S0950268820001235.

17. Z. Gao, Y. Xu, C. Sun, X. Wang, Y. Guo, S. Qiu, K. Ma, A systematic review of asymptomatic infections with COVID-19 J. Microbiol. Immunol. Infect. (2021), doi:10.1016/j.jmii.2020.05.001.

18. M. C. Rowbotham, K. L. Petersen, Zoster-associated pain and neural dysfunction Pain (2001), doi:10.1016/S0304-3959(01)00328-1.

19. S. Verma, L. Estanislao, D. Simpson, HIV-associated neuropathic pain: Epidemiology, pathophysiology and management. CNS Drugs 19, 325–334 (2005).

20. Y. Wu, X. Xu, Z. Chen, J. Duan, K. Hashimoto, L. Yang, C. Liu, C. Yang, Nervous system involvement after infection with COVID-19 and other coronaviruses Brain. Behav. Immun. (2020), doi:10.1016/j.bbi.2020.03.031.

21. E. M. Garry, A. Delaney, H. A. Anderson, E. C. Sirinathsinghji, R. H. Clapp, W. J. Martin, P. R. Kinchington, D. L. Krah, C. Abbadie, S. M. Fleetwood-Walker, Varicella zoster virus induces neuropathic changes in rat dorsal root ganglia and behavioral reflex sensitisation that is attenuated by gabapentin or sodium channel blocking drugs. Pain (2005), doi:10.1016/j.pain.2005.08.003.

22. I. Takasaki, T. Andoh, K. Shiraki, Y. Kuraishi, Allodynia and hyperalgesia induced by herpes simplex virus type-1 infection in mice. Pain (2000), doi:10.1016/S0304-3959(00)00240-2.

23. C. A. Pardo, J. C. McArthur, J. W. Griffin, in Journal of the Peripheral Nervous System, (2001).

24. K. Maratou, V. C. J. Wallace, F. S. Hasnie, K. Okuse, R. Hosseini, N. Jina, J. Blackbeard, T. Pheby, C. Orengo, A. H. Dickenson, S. B. McMahon, A. S. C. Rice, Comparison of dorsal root ganglion gene expression in rat models of traumatic and HIV-associated neuropathic pain. Eur. J. Pain (2009), doi:10.1016/j.ejpain.2008.05.011.

25. A. L. Oaklander, Clinical significance of angiotensin-converting enzyme 2 receptors for severe acute respiratory syndrome coronavirus 2 (COVID-19) on peripheral small-fiber sensory neurons is unknown today Pain (2020), doi:10.1097/j.pain.0000000000002050.

26. A. Paniz-Mondolfi, C. Bryce, Z. Grimes, R. E. Gordon, J. Reidy, J. Lednicky, E. M. Sordillo, M. Fowkes, Central nervous system involvement by severe acute respiratory syndrome coronavirus-2 (SARS-CoV-2) J. Med. Virol. (2020), doi:10.1002/jmv.25915.

27. J. Matschke, M. Lütgehetmann, C. Hagel, J. P. Sperhake, A. S. Schröder, C. Edler, H. Mushumba, A. Fitzek, L. Allweiss, M. Dandri, M. Dottermusch, A. Heinemann, S. Pfefferle, M. Schwabenland, D. Sumner Magruder, S. Bonn, M. Prinz, C. Gerloff, K. Püschel, S. Krasemann, M. Aepfelbacher, M. Glatzel, Neuropathology of patients with COVID-19 in Germany: a post-mortem case series. Lancet Neurol. 19 (2020), doi:10.1016/S1474-4422(20)30308-2.

28. B. Schurink, E. Roos, T. Radonic, E. Barbe, C. S. C. Bouman, H. H. de Boer, G. J. de Bree, E. B. Bulle, E. M. Aronica, S. Florquin, J. Fronczek, L. M. A. Heunks, M. D. de Jong, L. Guo, R. du Long, R. Lutter, P. C. G. Molenaar, E. A. Neefjes-Borst, H. W. M. Niessen, C. J. M. van Noesel, J. J. T. H. Roelofs, E. J. Snijder, E. C. Soer, J. Verheij, A. P. J. Vlaar, W. Vos, N. N. van der Wel, A. C. van der Wal, P. van der Valk, M. Bugiani, Viral presence and immunopathology in patients with lethal COVID-19: a prospective autopsy cohort study. The Lancet Microbe 1 (2020), doi:10.1016/S2666-5247(20)30144-0.

29. K. T. Thakur, E. H. Miller, M. D. Glendinning, O. Al-Dalahmah, M. A. Banu, A. K. Boehme, A. L. Boubour, S. S. Bruce, A. M. Chong, J. Claassen, P. L. Faust, G. Hargus, R. A. Hickman, S. Jambawalikar, A. G. Khandji, C. Y. Kim, R. S. Klein, A. Lignelli-Dipple, C. C. Lin, Y. Liu, M. L. Miller, G. Moonis, A. S. Nordvig, J. B. Overdevest, M. L. Prust, S. Przedborski, W. H. Roth, A. Soung, K. Tanji, A. F. Teich, D. Agalliu, A. C. Uhlemann, J. E. Goldman, P. Canoll, COVID-19 neuropathology at Columbia University Irving Medical Center/New York Presbyterian Hospital. Brain 144 (2021), doi:10.1093/brain/awab148.

30. T. Menter, J. D. Haslbauer, R. Nienhold, S. Savic, H. Hopfer, N. Deigendesch, S. Frank, D. Turek, N. Willi, H. Pargger, S. Bassetti, J. D. Leuppi, G. Cathomas, M. Tolnay, K. D. Mertz, A. Tzankov, Postmortem examination of COVID-19 patients reveals diffuse alveolar damage with severe capillary congestion and variegated findings in lungs and other organs suggesting vascular dysfunction. Histopathology 77 (2020), doi:10.1111/his.14134.

31. D. A. Hoagland, R. Møller, S. A. Uhl, K. Oishi, J. Frere, I. Golynker, S. Horiuchi, M. Panis, D. Blanco-Melo, D. Sachs, K. Arkun, J. K. Lim, B. R. tenOever, Leveraging the antiviral type I interferon system as a first line of defense against SARS-CoV-2 pathogenicity. Immunity 54 (2021), doi:10.1016/j.immuni.2021.01.017.

32. J. J. Frere, R. A. Serafini, K. D. Pryce, M. Zazhytska, K. Oishi, I. Golynker, M. Panis, J. Zimering, S. Horiuchi, D. A. Hoagland, R. Møller, A. Ruiz, A. Kodra, J. B. Overdevest, P. D. Canoll, A. C. Borczuk, V. Chandar, Y. Bram, R. Schwartz, S. Lomvardas, V. Zachariou, B. R. TenOever, SARS-CoV-2 infection in hamsters and humans results in lasting and unique systemic perturbations post recovery. Sci. Transl. Med. (2022) (available at https://www.science.org/doi/full/10.1126/scitranslmed.abq3059).

33. L. Zanin, G. Saraceno, P. P. Panciani, G. Renisi, L. Signorini, K. Migliorati, M. M. Fontanella, SARS-CoV-2 can induce brain and spine demyelinating lesions. Acta Neurochir. (Wien). (2020), doi:10.1007/s00701-020-04374-x.

34. M. F. V. V. Aragão, M. C. Leal, O. Q. Cartaxo Filho, T. M. Fonseca, M. M. Valença, Anosmia in COVID-19 associated with injury to the olfactory bulbs evident on MRI. Am. J. Neuroradiol. (2020), doi:10.3174/ajnr.A6675.

35. F. Roy-Gash, D. M. Marine, D. Jean-Michel, V. Herve, B. Raphael, E. Nicolas, COVID-19-associated acute cerebral venous thrombosis: Clinical, CT, MRI and EEG features Crit. Care (2020), doi:10.1186/s13054-020-03131-x.

36. E. Carroll, A. Lewis, Catastrophic Intracranial Hemorrhage in Two Critically Ill Patients with COVID-19. Neurocrit. Care (2020), doi:10.1007/s12028-020-00993-5.

37. C. E. Fernandez, C. K. Franz, J. H. Ko, J. M. Walter, I. J. Koralnik, S. Ahlawat, S. Deshmukh, Imaging review of peripheral nerve injuries in patients with COVID-19Radiology (2021), doi:10.1148/radiol.2020203116.

38. R. M. C. Abrams, D. M. Simpson, A. Navis, N. Jette, L. Zhou, S. C. Shin, Small fiber neuropathy associated with SARS-CoV-2 infection. Muscle and Nerve (2021), doi:10.1002/mus.27458.

39. B. Neumann, M. L. Schmidbauer, K. Dimitriadis, S. Otto, B. Knier, W. D. Niesen, J. A. Hosp, A. Günther, S. Lindemann, G. Nagy, T. Steinberg, R. A. Linker, B. Hemmer, J. Bösel, Cerebrospinal fluid findings in COVID-19 patients with neurological symptoms J. Neurol. Sci. (2020), doi:10.1016/j.jns.2020.117090.

40. G. Destras, A. Bal, V. Escuret, F. Morfin, B. Lina, L. Josset, Systematic SARS-CoV-2 screening in cerebrospinal fluid during the COVID-19 pandemic The Lancet Microbe (2020), doi:10.1016/S2666-5247(20)30066-5.

41. Y. H. Huang, D. Jiang, J. T. Huang, SARS-CoV-2 Detected in Cerebrospinal Fluid by PCR in a Case of COVID-19 Encephalitis Brain. Behav. Immun. (2020), doi:10.1016/j.bbi.2020.05.012.

42. R. B. Domingues, M. C. Mendes-Correa, F. B. V. de Moura Leite, E. C. Sabino, D. Z. Salarini, I. Claro, D. W. Santos, J. G. de Jesus, N. E. Ferreira, C. M. Romano, C. A. S. Soares, First case of SARS-COV-2 sequencing in cerebrospinal fluid of a patient with suspected demyelinating disease J. Neurol. (2020), doi:10.1007/s00415-020-09996-w.

43. Y. C. Li, W. Z. Bai, N. Hirano, T. Hayashida, T. Hashikawa, Coronavirus infection of rat dorsal root ganglia: Ultrastructural characterization of viral replication, transfer, and the early response of satellite cells. Virus Res. (2012), doi:10.1016/j.virusres.2011.12.021.

44. S. Perlman, G. Jacobsen, A. L. Olson, A. Afifi, Identification of the spinal cord as a major site of persistence during during chronic infection with a murine coronavirus. Virology (1990), doi:10.1016/0042-6822(90)90426-R.

45. R. Elliott, F. Li, I. Dragomir, M. M. W. Chua, B. D. Gregory, S. R. Weiss, Analysis of the Host Transcriptome from Demyelinating Spinal Cord of Murine Coronavirus-Infected Mice. PLoS One (2013), doi:10.1371/journal.pone.0075346.

46. O. O. Koyuncu, I. B. Hogue, L. W. Enquist, Virus infections in the nervous system Cell Host Microbe (2013), doi:10.1016/j.chom.2013.03.010.

47. S. Hosseini, E. Wilk, K. Michaelsen-Preusse, I. Gerhauser, W. Baumgärtner, R. Geffers, K. Schughart, M. Korte, Long-term neuroinflammation induced by influenza a virus infection and the impact on hippocampal neuron morphology and function. J. Neurosci. (2018), doi:10.1523/JNEUROSCI.1740-17.2018.

48. K. Dobrindt, D. A. Hoagland, C. Seah, B. Kassim, C. P. O’Shea, A. Murphy, M. Iskhakova, M. B. Fernando, S. K. Powell, P. J. M. Deans, B. Javidfar, C. Peter, R. Møller, S. A. Uhl, M. F. Garcia, M. Kimura, K. Iwasawa, J. F. Crary, D. N. Kotton, T. Takebe, L. M. Huckins, B. R. tenOever, S. Akbarian, K. J. Brennand, Common Genetic Variation in Humans Impacts In Vitro Susceptibility to SARS-CoV-2 Infection. Stem Cell Reports 16 (2021), doi:10.1016/j.stemcr.2021.02.010.

49. E. Song, C. Zhang, B. Israelow, A. Lu-Culligan, A. V. Prado, S. Skriabine, P. Lu, O. El Weizman, F. Liu, Y. Dai, K. Szigeti-Buck, Y. Yasumoto, G. Wang, C. Castaldi, J. Heltke, E. Ng, J. Wheeler, M. M. Alfajaro, E. Levavasseur, B. Fontes, N. G. Ravindra, D. van Dijk, S. Mane, M. Gunel, A. Ring, S. A. Jaffar Kazmi, K. Zhang, C. B. Wilen, T. L. Horvath, I. Plu, S. Haik, J. L. Thomas, A. Louvi, S. F. Farhadian, A. Huttner, D. Seilhean, N. Renier, K. Bilguvar, A. Iwasaki, Neuroinvasion of SARS-CoV-2 in human and mouse brain. J. Exp. Med. 218 (2021), doi:10.1084/JEM.20202135.

50. S. F. Sia, L. M. Yan, A. W. H. Chin, K. Fung, K. T. Choy, A. Y. L. Wong, P. Kaewpreedee, R. A. P. M. Perera, L. L. M. Poon, J. M. Nicholls, M. Peiris, H. L. Yen, Pathogenesis and transmission of SARS-CoV-2 in golden hamsters. Nature 583 (2020), doi:10.1038/s41586-020-2342-5.

51. D. J. Morales, D. J. Lenschow, The antiviral activities of ISG15 J. Mol. Biol. 425 (2013), doi:10.1016/j.jmb.2013.09.041.

52. P. Barragán-Iglesias, Ú. Franco-Enzástiga, V. Jeevakumar, S. Shiers, A. Wangzhou, V. Granados-Soto, Z. T. Campbell, G. Dussor, T. J. Price, Type I interferons act directly on nociceptors to produce pain sensitization: Implications for viral infection-induced pain. J. Neurosci. 40 (2020), doi:10.1523/JNEUROSCI.3055-19.2020.

53. R. Eccles, Understanding the symptoms of the common cold and influenza Lancet Infect. Dis. (2005), doi:10.1016/S1473-3099(05)70270-X.

54. M. Pietzner, E. Wheeler, J. Carrasco-Zanini, J. Raffler, N. D. Kerrison, E. Oerton, V. P. W. Auyeung, J. Luan, C. Finan, J. P. Casas, R. Ostroff, S. A. Williams, G. Kastenmüller, M. Ralser, E. R. Gamazon, N. J. Wareham, A. D. Hingorani, C. Langenberg, Genetic architecture of host proteins involved in SARS-CoV-2 infection. Nat. Commun. 11 (2020), doi:10.1038/s41467-020-19996-z.

55. A. Sharma, M. A. Pollett, G. W. Plant, A. R. Harvey, Changes in mRNA expression of class 3 semaphorins and their receptors in the adult rat retino-collicular system after unilateral optic nerve injury. Investig. Ophthalmol. Vis. Sci. (2012), doi:10.1167/iovs.12-10799.

56. A. Verheyen, E. Peeraer, R. Nuydens, J. Dhondt, K. Poesen, I. Pintelon, A. Daniels, J. P. Timmermans, T. Meert, P. Carmeliet, D. Lambrechts, Systemic anti-vascular endothelial growth factor therapies induce a painful sensory neuropathy. Brain (2012), doi:10.1093/brain/aws145.

57. A. Moutal, L. F. Martin, L. Boinon, K. Gomez, D. Ran, Y. Zhou, H. J. Stratton, S. Cai, S. Luo, K. B. Gonzalez, S. Perez-Miller, A. Patwardhan, M. M. Ibrahim, R. Khanna, SARS-CoV-2 spike protein co-opts VEGF-A/neuropilin-1 receptor signaling to induce analgesia. Pain (2021), doi:10.1097/j.pain.0000000000002097.

58. G. Taccola, P. J. Doyen, J. Damblon, N. Dingu, B. Ballarin, A. Steyaert, A. des Rieux, P. Forget, E. Hermans, B. Bosier, R. Deumens, A new model of nerve injury in the rat reveals a role of Regulator of G protein Signaling 4 in tactile hypersensitivity. Exp. Neurol. (2016), doi:10.1016/j.expneurol.2016.09.008.

59. K. Prasad, F. Khatoon, S. Rashid, N. Ali, A. F. AlAsmari, M. Z. Ahmed, A. S. Alqahtani, M. S. Alqahtani, V. Kumar, Targeting hub genes and pathways of innate immune response in COVID-19: A network biology perspective. Int. J. Biol. Macromol. (2020), doi:10.1016/j.ijbiomac.2020.06.228.

60. L. Provenzi, M. Fumagalli, G. Scotto di Minico, R. Giorda, F. Morandi, I. Sirgiovanni, P. Schiavolin, F. Mosca, R. Borgatti, R. Montirosso, Pain-related increase in serotonin transporter gene methylation associates with emotional regulation in 4.5-year-old preterm-born children. Acta Paediatr. Int. J. Paediatr. (2020), doi:10.1111/apa.15077.

61. H. B. Raju, N. F. Tsinoremas, E. Capobianco, Emerging putative associations between non-coding RNAs and protein-coding genes in neuropathic pain: Added value from reusing microarray data. Front. Neurol. (2016), doi:10.3389/fneur.2016.00168.

62. X. Li, W. Wang, Q. Chen, Y. Zhou, L. Wang, H. Huang, Antinociceptive effects of IL-6R vs. glucocorticoid receptors during rat hind paw inflammatory pain. Neurosci. Lett. (2020), doi:10.1016/j.neulet.2020.135356.

63. D. Zhang, J. Y. Mou, F. Wang, J. Liu, X. Hu, CRNDE enhances neuropathic pain via modulating miR-136/IL6R axis in CCI rat models. J. Cell. Physiol. (2019), doi:10.1002/jcp.28790.

64. H. Wakabayashi, S. Kato, N. Nagao, G. Miyamura, Y. Naito, A. Sudo, Interleukin-6 Inhibitor Suppresses Hyperalgesia Without Improvement in Osteoporosis in a Mouse Pain Model of Osteoporosis. Calcif. Tissue Int. (2019), doi:10.1007/s00223-019-00521-4.

65. R. R. Ji, H. Baba, G. J. Brenner, C. J. Woolf, Nociceptive-specific activation of ERK in spinal neurons contributes to pain hypersensitivity. Nat. Neurosci. (1999), doi:10.1038/16040.

66. A. Ciruela, A. K. Dixon, S. Bramwell, M. I. Gonzalez, R. D. Pinnock, K. Lee, Identification of MEK1 as a novel target for the treatment of neuropathic pain. Br. J. Pharmacol. (2003), doi:10.1038/sj.bjp.0705103.

67. S. Yamakita, Y. Horii, H. Takemura, Y. Matsuoka, A. Yamashita, Y. Yamaguchi, M. Matsuda, T. Sawa, F. Amaya, Synergistic activation of ERK1/2 between A-fiber neurons and glial cells in the DRG contributes to pain hypersensitivity after tissue injury. Mol. Pain (2018), doi:10.1177/1744806918767508.

68. J. Zhang, J. Jiang, G. Bao, G. Xu, L. Wang, J. Chen, C. Chen, C. Wu, P. Xue, D. Xu, Y. Sun, Z. Cui, Interaction between C/EBPβ and RUNX2 promotes apoptosis of chondrocytes during human lumbar facet joint degeneration. J. Mol. Histol. (2020), doi:10.1007/s10735-020-09891-8.

69. S. J. Rice, G. Aubourg, A. K. Sorial, D. Almarza, M. Tselepi, D. J. Deehan, L. N. Reynard, J. Loughlin, Identification of a novel, methylation-dependent, RUNX2 regulatory region associated with osteoarthritis risk. Hum. Mol. Genet. (2018), doi:10.1093/hmg/ddy257.

70. A. Tajar, J. McBeth, D. M. Lee, G. J. MacFarlane, I. T. Huhtaniemi, J. D. Finn, G. Bartfai, S. Boonen, F. F. Casanueva, G. Forti, A. Giwercman, T. S. Han, K. Kula, F. Labrie, M. E. J. Lean, N. Pendleton, M. Punab, A. J. Silman, D. Vanderschueren, T. W. O’Neill, F. C. W. Wu, Elevated levels of gonadotrophins but not sex steroids are associated with musculoskeletal pain in middle-aged and older European men. Pain (2011), doi:10.1016/j.pain.2011.01.048.

71. M. X. Xie, X. Y. Cao, W. A. Zeng, R. C. Lai, L. Guo, J. C. Wang, Y. Bin Xiao, X. Zhang, D. Chen, X. G. Liu, X. L. Zhang, ATF4 selectively regulates heat nociception and contributes to kinesin-mediated TRPM3 trafficking. Nat. Commun. (2021), doi:10.1038/s41467-021-21731-1.

72. L. Dong, B. B. Guarino, K. L. Jordan-Sciutto, B. A. Winkelstein, Activating transcription factor 4, a mediator of the integrated stress response, is increased in the dorsal root ganglia following painful facet joint distraction. Neuroscience (2011), doi:10.1016/j.neuroscience.2011.07.059.

73. O. Q. Russe, C. V. Möser, K. L. Kynast, T. S. King, H. Stephan, G. Geisslinger, E. Niederberger, Activation of the AMP-activated protein kinase reduces inflammatory nociception. J. Pain (2013), doi:10.1016/j.jpain.2013.05.012.

74. T. S. King-Himmelreich, C. V. Möser, M. C. Wolters, J. Schmetzer, Y. Schreiber, N. Ferreirós, O. Q. Russe, G. Geisslinger, E. Niederberger, AMPK contributes to aerobic exercise-induced antinociception downstream of endocannabinoids. Neuropharmacology (2017), doi:10.1016/j.neuropharm.2017.05.002.

75. G. Giaccone, P. Zatloukal, J. Roubec, K. Floor, J. Musil, M. Kuta, R. J. Van Klaveren, S. Chaudhary, A. Gunther, S. Shamsili, Multicenter phase II trial of YM155, a small-molecule suppressor of survivin, in patients with advanced, refractory, non-small-cell lung cancer. J. Clin. Oncol. (2009), doi:10.1200/JCO.2008.21.1862.

76. M. R. Clemens, O. A. Gladkov, E. Gartner, V. Vladimirov, J. Crown, J. Steinberg, F. Jie, A. Keating, Phase II, multicenter, open-label, randomized study of YM155 plus docetaxel as first-line treatment in patients with HER2-negative metastatic breast cancer. Breast Cancer Res. Treat. (2015), doi:10.1007/s10549-014-3238-6.

77. A. W. Tolcher, D. I. Quinn, A. Ferrari, F. Ahmann, G. Giaccone, T. Drake, A. Keating, J. S. De Bono, A phase II study of YM155, a novel small-molecule suppressor of survivin, in castration-resistant taxane-pretreated prostate cancer. Ann. Oncol. (2012), doi:10.1093/annonc/mdr353.

78. T. Yamauchi, N. Nakamura, M. Hiramoto, M. Yuri, H. Yokota, M. Naitou, M. Takeuchi, K. Yamanaka, A. Kita, T. Nakahara, I. Kinoyama, A. Matsuhisa, N. Kaneko, H. Koutoku, M. Sasamata, M. Kobori, M. Katou, S. Tawara, S. Kawabata, K. Furuichi, Sepantronium Bromide (YM155) induces disruption of the ILF3/p54nrb complex, which is required for survivin expression. Biochem. Biophys. Res. Commun. (2012), doi:10.1016/j.bbrc.2012.07.103.

79. T. Nakahara, M. Takeuchi, I. Kinoyama, T. Minematsu, K. Shirasuna, A. Matsuhisa, A. Kita, F. Tominaga, K. Yamanaka, M. Kudoh, M. Sasamata, YM155, a novel small-molecule survivin suppressant, induces regression of established human hormone-refractory prostate tumor xenografts. Cancer Res. (2007), doi:10.1158/0008-5472.CAN-07-1343.

80. K. Avrampou, K. D. Pryce, A. Ramakrishnan, F. Sakloth, S. Gaspari, R. A. Serafini, V. Mitsi, C. Polizu, C. Swartz, B. Ligas, A. Richards, L. Shen, F. B. Carr, V. Zachariou, RGS4 Maintains Chronic Pain Symptoms in Rodent Models. J. Neurosci. 39, 8291–8304 (2019).

81. E. Vachon-Presseau, P. Tétreault, B. Petre, L. Huang, S. E. Berger, S. Torbey, A. T. Baria, A. R. Mansour, J. A. Hashmi, J. W. Griffith, E. Comasco, T. J. Schnitzer, M. N. Baliki, A. V. Apkarian, Corticolimbic anatomical characteristics predetermine risk for chronic pain. Brain 139, 1958–1970 (2016).

82. P. Muglia, F. Tozzi, N. W. Galwey, C. Francks, R. Upmanyu, X. Q. Kong, A. Antoniades, E. Domenici, J. Perry, S. Rothen, C. L. Vandeleur, V. Mooser, G. Waeber, P. Vollenweider, M. Preisig, S. Lucae, B. Müller-Myhsok, F. Holsboer, L. T. Middleton, A. D. Roses, Genome-wide association study of recurrent major depressive disorder in two European case-control cohorts. Mol. Psychiatry 15 (2010), doi:10.1038/mp.2008.131.

83. Y. Wang, L. Li, C. Xu, X. Cao, Z. Liu, N. Sun, A. Zhang, X. Li, K. Zhang, Polymorphism of ERK/PTPRR Genes in Major Depressive Disorder at Resting-State Brain Function. Dev. Neuropsychol. 42 (2017), doi:10.1080/87565641.2017.1306527.

84. M. Muriello, A. Y. Kim, K. Sondergaard Schatz, N. Beck, M. Gunay-Aygun, J. E. Hoover-Fong, Growth hormone deficiency, aortic dilation, and neurocognitive issues in Feingold syndrome 2. Am. J. Med. Genet. Part A 179 (2019), doi:10.1002/ajmg.a.61037.

85. H. Ganjavi, V. M. Siu, M. Speevak, P. A. MacDonald, A fourth case of Feingold syndrome type 2: Psychiatric presentation and management. BMJ Case Rep. 2014 (2014), doi:10.1136/bcr-2014-207501.

86. Y. Wen, X. Fan, H. Bu, L. Ma, C. Kong, C. Huang, Y. Xu, Downregulation of lncRNA FIRRE relieved the neuropathic pain of female mice by suppressing HMGB1 expression. Mol. Cell. Biochem. 476 (2021), doi:10.1007/s11010-020-03949-7.

87. K. Gómez, A. Sandoval, P. Barragán-Iglesias, V. Granados-Soto, R. Delgado-Lezama, R. Felix, R. González-Ramírez, Transcription Factor Sp1 Regulates the Expression of Calcium Channel α2δ-1 Subunit in Neuropathic Pain. Neuroscience (2019), doi:10.1016/j.neuroscience.2019.06.011.

88. S. Li, F. Zhao, Q. Tang, C. Xi, J. He, Y. Wang, M. X. Zhu, Z. Cao, Sarco/endoplasmic reticulum Ca 2+ -ATPase 2b mediates oxidation-induced endoplasmic reticulum stress to regulate neuropathic pain. Br. J. Pharmacol. (2021), doi:10.1111/bph.15744.

89. D. J. Kao, A. H. Li, J. C. Chen, R. S. Luo, Y. L. Chen, J. C. Lu, H. L. Wang, CC chemokine ligand 2 upregulates the current density and expression of TRPV1 channels and Nav1.8 sodium channels in dorsal root ganglion neurons. J. Neuroinflammation (2012), doi:10.1186/1742-2094-9-189.

90. W. Xie, Z. Y. Tan, C. Barbosa, J. A. Strong, T. R. Cummins, J. M. Zhang, Upregulation of the sodium channel Na v ß4 subunit and its contributions to mechanical hypersensitivity and neuronal hyperexcitability in a rat model of radicular pain induced by local dorsal root ganglion inflammation. Pain (2016), doi:10.1097/j.pain.0000000000000453.

91. C. Barbosa, Z. Y. Tan, R. Wang, W. Xie, J. A. Strong, R. R. Patel, M. R. Vasko, J. M. Zhang, T. R. Cummins, Navβ4 regulates fast resurgent sodium currents and excitability in sensory neurons. Mol. Pain (2015), doi:10.1186/s12990-015-0063-9.

92. A. Broyl, S. L. Corthals, J. L. M. Jongen, B. van der Holt, R. Kuiper, Y. de Knegt, M. van Duin, L. el Jarari, U. Bertsch, H. M. Lokhorst, B. G. Durie, H. Goldschmidt, P. Sonneveld, Mechanisms of peripheral neuropathy associated with bortezomib and vincristine in patients with newly diagnosed multiple myeloma: a prospective analysis of data from the HOVON-65/GMMG-HD4 trial. Lancet Oncol. (2010), doi:10.1016/S1470-2045(10)70206-0.

93. N. Munawar, M. A. Oriowo, W. Masocha, Antihyperalgesic activities of endocannabinoids in a mouse model of antiretroviral-Induced neuropathic pain. Front. Pharmacol. (2017), doi:10.3389/fphar.2017.00136.

94. R. Fu, Y. Tang, W. Li, Z. Ren, D. Li, J. Zheng, W. Zuo, X. Chen, Q. K. Zuo, K. L. Tam, Y. Zou, T. Bachmann, A. Bekker, J. H. Ye, Endocannabinoid signaling in the lateral habenula regulates pain and alcohol consumption. Transl. Psychiatry (2021), doi:10.1038/s41398-021-01337-3.

95. Z. Hu, N. Deng, K. Liu, N. Zhou, Y. Sun, W. Zeng, CNTF-STAT3-IL-6 Axis Mediates Neuroinflammatory Cascade across Schwann Cell-Neuron-Microglia. Cell Rep. (2020), doi:10.1016/j.celrep.2020.107657.

96. H. J. Chung, J. D. Kim, K. H. Kim, N. Y. Jeong, G protein-coupled receptor, family C, group 5 (GPRC5B) downregulation in spinal cord neurons is involved in neuropathic pain. Korean J. Anesthesiol. (2014), doi:10.4097/kjae.2014.66.3.230.

97. L. F. Ferrari, O. Bogen, N. Alessandri-Haber, E. Levine, R. W. Gear, J. D. Levine, Transient decrease in nociceptor GRK2 expression produces long-term enhancement in inflammatory pain. Neuroscience (2012), doi:10.1016/j.neuroscience.2012.07.004.

98. N. Eijkelkamp, C. J. Heijnen, H. L. D. M. Willemen, R. Deumens, E. A. J. Joosten, W. Kleibeuker, I. J. M. Den Hartog, C. T. J. Van Velthoven, C. Nijboer, M. A. Nassar, G. W. Dorn, J. N. Wood, A. Kavelaars, GRK2: A novel cell-specific regulator of severity and duration of inflammatory pain. J. Neurosci. (2010), doi:10.1523/JNEUROSCI.5752-09.2010.

99. M. Parisien, A. Samoshkin, S. N. Tansley, M. H. Piltonen, L. J. Martin, N. El-Hachem, C. Dagostino, M. Allegri, J. S. Mogil, A. Khoutorsky, L. Diatchenko, Genetic pathway analysis reveals a major role for extracellular matrix organization in inflammatory and neuropathic pain. Pain (2019), doi:10.1097/j.pain.0000000000001471.

100. M. Imai, K. Iwatsuki-Horimoto, M. Hatta, S. Loeber, P. J. Halfmann, N. Nakajima, T. Watanabe, M. Ujie, K. Takahashi, M. Ito, S. Yamada, S. Fan, S. Chiba, M. Kuroda, L. Guan, K. Takada, T. Armbrust, A. Balogh, Y. Furusawa, M. Okuda, H. Ueki, A. Yasuhara, Y. Sakai-Tagawa, T. J. S. Lopes, M. Kiso, S. Yamayoshi, N. Kinoshita, N. Ohmagari, S. I. Hattori, M. Takeda, H. Mitsuya, F. Krammer, T. Suzuki, Y. Kawaoka, Syrian hamsters as a small animal model for SARS-CoV-2 infection and countermeasure development. Proc. Natl. Acad. Sci. U. S. A. 117 (2020), doi:10.1073/pnas.2009799117.

101. C. Muñoz-Fontela, W. E. Dowling, S. G. P. Funnell, P. S. Gsell, A. X. Riveros-Balta, R. A. Albrecht, H. Andersen, R. S. Baric, M. W. Carroll, M. Cavaleri, C. Qin, I. Crozier, K. Dallmeier, L. de Waal, E. de Wit, L. Delang, E. Dohm, W. P. Duprex, D. Falzarano, C. L. Finch, M. B. Frieman, B. S. Graham, L. E. Gralinski, K. Guilfoyle, B. L. Haagmans, G. A. Hamilton, A. L. Hartman, S. Herfst, S. J. F. Kaptein, W. B. Klimstra, I. Knezevic, P. R. Krause, J. H. Kuhn, R. Le Grand, M. G. Lewis, W. C. Liu, P. Maisonnasse, A. K. McElroy, V. Munster, N. Oreshkova, A. L. Rasmussen, J. Rocha-Pereira, B. Rockx, E. Rodríguez, T. F. Rogers, F. J. Salguero, M. Schotsaert, K. J. Stittelaar, H. J. Thibaut, C. Te Tseng, J. Vergara-Alert, M. Beer, T. Brasel, J. F. W. Chan, A. García-Sastre, J. Neyts, S. Perlman, D. S. Reed, J. A. Richt, C. J. Roy, J. Segalés, S. S. Vasan, A. M. Henao-Restrepo, D. H. Barouch, Animal models for COVID-19 Nature 586 (2020), doi:10.1038/s41586-020-2787-6.

102. G. Gao, W. Li, S. Liu, D. Han, X. Yao, J. Jin, D. Han, W. Sun, X. Chen, The positive feedback loop between ILF3 and lncrna ILF3-AS1 promotes melanoma proliferation, migration, and invasion. Cancer Manag. Res. (2018), doi:10.2147/CMAR.S186777.

103. X. Yang, F. Lin, F. Gao, Up-regulated long non-coding RNA ILF3-AS1 indicates poor prognosis of nasopharyngeal carcinoma and promoted cell metastasis. Int. J. Biol. Markers (2020), doi:10.1177/1724600820955199.

104. X. hui Hu, J. Dai, H. lai Shang, Z. xue Zhao, Y. dong Hao, SP1-mediated upregulation of lncRNA ILF3-AS1 functions a ceRNA for miR-212 to contribute to osteosarcoma progression via modulation of SOX5. Biochem. Biophys. Res. Commun. (2019), doi:10.1016/j.bbrc.2019.02.110.

105. M. D. Sanna, N. Galeotti, The HDAC1/c-JUN complex is essential in the promotion of nerve injury-induced neuropathic pain through JNK signaling. Eur. J. Pharmacol. 825 (2018), doi:10.1016/j.ejphar.2018.02.034.

106. F. Sakloth, L. Manouras, K. Avrampou, V. Mitsi, R. A. Serafini, K. D. Pryce, V. Cogliani, O. Berton, M. Jarpe, V. Zachariou, HDAC6-selective inhibitors decrease nerve-injury and inflammation-associated mechanical hypersensitivity in mice. Psychopharmacology (Berl). 237 (2020), doi:10.1007/s00213-020-05525-9.

107. I. Vasileiou, I. Adamakis, E. Patsouris, S. Theocharis, Ephrins and painExpert Opin. Ther. Targets (2013), doi:10.1517/14728222.2013.801456.

108. A. Leroux, B. Paiva dos Santos, J. Leng, H. Oliveira, J. Amédée, Sensory neurons from dorsal root ganglia regulate endothelial cell function in extracellular matrix remodelling. Cell Commun. Signal. (2020), doi:10.1186/s12964-020-00656-0.

109. M. E. Kerrisk, L. A. Cingolani, A. J. Koleske, in Progress in Brain Research, (2014).

110. N. N. Knezevic, A. Yekkirala, T. L. Yaksh, Basic/Translational Development of Forthcoming Opioid- and Nonopioid-Targeted Pain Therapeutics. Anesth. Analg. 125, 1714–1732 (2017).

111. K. D. Pryce, H. J. Kang, F. Sakloth, Y. Liu, S. Khan, K. Toth, A. Kapoor, A. Nicolais, T. Che, L. Qin, F. Bertherat, H. Ü. Kaniskan, J. Jin, M. D. Cameron, B. L. Roth, V. Zachariou, M. Filizola, A promising chemical series of positive allosteric modulators of the μ-opioid receptor that enhance the antinociceptive efficacy of opioids but not their adverse effects. Neuropharmacology 195 (2021), doi:10.1016/j.neuropharm.2021.108673.

112. M. I. Love, W. Huber, S. Anders, Moderated estimation of fold change and dispersion for RNA-seq data with DESeq2. Genome Biol. (2014), doi:10.1186/s13059-014-0550-8.

113. E. Y. Chen, C. M. Tan, Y. Kou, Q. Duan, Z. Wang, G. V. Meirelles, N. R. Clark, A. Ma’ayan, Enrichr: Interactive and collaborative HTML5 gene list enrichment analysis tool. BMC Bioinformatics (2013), doi:10.1186/1471-2105-14-128.

